# Resting-state alpha and mu rhythms change shape across development but lack diagnostic sensitivity for ADHD and autism

**DOI:** 10.1101/2023.10.13.562301

**Authors:** Andrew Bender, Bradley Voytek, Natalie Schaworonkow

**Author notes:** contributed equally to the work (Bradley Voytek, Natalie Schaworonkow).

## Abstract

In the human brain, the alpha rhythm in occipital cortex and the mu rhythm in sensorimotor cortex are among the most prominent rhythms, with both rhythms functionally implicated in gating modality-specific information. Separation of these rhythms is non-trivial due to the spatial mixing of these oscillations in sensor space. Using a computationally efficient processing pipeline requiring no manual data cleaning, we isolated alpha and/or mu rhythms from electroencephalography recordings performed on 1605 children aged 5–18. Using the extracted time series for each rhythm, we characterized the waveform shape on a cycle-by-cycle basis and examined whether and how the waveform shape differs across development. We demonstrate that alpha and mu rhythms both exhibit nonsinusoidal waveform shape that changes significantly across development, in addition to the known large changes in oscillatory frequency. This dataset also provided an opportunity to assess oscillatory measures for attention-deficit hyperactivity disorder (ADHD) and autism spectrum disorder (ASD). We found no differences in the resting-state features of these alpha-band rhythms for either ADHD or ASD in comparison to typically developing participants in this dataset. While waveform shape is ignored by traditional Fourier spectral analyses, these nonsinusoidal properties may be informative for building more constrained generative models for different types of alpha-band rhythms, yielding more specific insight into their generation.

## Introduction

Rhythms in the alpha band (8–13 Hz) are the most prominent rhythms in the human brain (Klimesch, 2012). The occipital alpha rhythm is most noticeable during wakeful periods when participants’ eyes are closed (Berger, 1929), but also changes dynamically during attentional tasks, decreasing in brain areas related to task-specific visual input (Kelly et al., 2006; Thut et al., 2006; Worden et al., 2000). By contrast, the sensorimotor mu rhythm is strongest when a participant is not moving (Salmelin and Hari, 1994). Mu activity changes dynamically during motor tasks, decreasing in power in motor areas controlling moving body parts and increasing in power in motor areas controlling immobile body parts (Pfurtscheller and Neuper, 1994). In this way, both alpha and mu rhythms are important for gating modality-specific information, inhibiting the processing of functionally irrelevant information.

These rhythms share a frequency band and display a similar change in frequency across development, one of the strongest effects in the field of oscillations (Freschl et al., 2022; Lindsley, 1939). Despite this commonality, the rhythms show distinct waveform shape: alpha rhythms appear triangular in raw traces (Stam et al., 1999), while mu rhythms are considered to have an arc-shaped waveform (HJ Gastaut and Bert, 1954; Kuhlman, 1978). The waveform shape likely reflects differences in the physiological properties of the underlying oscillatory generators (Cole and Voytek, 2017). Computational modeling has suggested that the sharpness of oscillatory peaks and troughs reflects the synchronicity of the underlying excitatory input currents (Sherman et al., 2016). The waveform shape of alpha-band rhythms has been characterized in EEG in sensor space (Schaworonkow and Nikulin, 2019) and MEG (Giehl and Siegel, 2024), but an investigation of waveform characteristics in developmental populations is lacking. With the main factor driving the prominent oscillatory frequency increase during development remaining unknown, such precise characterization will help determine whether the oscillatory changes are driven by gross anatomical changes or distinct modifications to the oscillatory generators underlying these rhythms.

Given the strong changes in oscillations across development and the link of waveform shape to pathophysiological properties of circuits in Parkinson’s disease (Cole, van der Meij, et al., 2017), we were interested in whether the waveform shape of resting-state alpha and mu rhythms acquired in a developmental population could serve as a biomarker. Alpha-band rhythms have been implicated in autism spectrum disorder (ASD) and attention deficit hyperactivity disorder (ADHD) in terms of both their frequency and power, though the results have been inconsistent (Freschl et al., 2022; Javitt et al., 2020). Our aim here is to repeat some of the typically performed analysis in previous literature comparing ASD and ADHD groups with participants without a diagnosis, with a robust sample size for frequency and amplitude measures, while also resolving between two different rhythm types and taking into account waveform shape.

The aim of the current study was therefore twofold: we investigated how waveform shape features of alpha and mu rhythms change both across development and neurodevelopmental disorders. Our study provides four methodological advantages over existing approaches. First, we analyze the signals primarily in the time domain (Cole and Voytek, 2019), to characterize nonsinusoidal features of these rhythms. Second, we use a large open developmental dataset (ages 5–18) of high-density, resting-state electroencephalography (EEG) (Alexander et al., 2017), allowing us to perform a high-powered comparison of alpha and mu rhythms across development. Third, we use spatio-spectral decomposition (SSD) (Nikulin, Nolte, and Curio, 2011), a statistical procedure using individual peak frequency information to identify different types of alpha-band rhythms that are mixed on the sensor level due to EEG volume conduction. This spatial mixing distorts waveform shape, making this spatial filtering step crucial (Schaworonkow and Nikulin, 2022). Finally, we use template-based source localization on these demixed alpha-band rhythms to isolate alpha and mu rhythms from other alpha-band rhythms, such as parietal alpha rhythms, which are prominent in resting-state EEG but outside the purview of the hypotheses tested herein.

We found that, while both alpha and mu are nonsinusoidal, mu rhythms have a significantly higher peak-trough asymmetry compared to alpha. Although alpha and mu rhythms are very similar in their frequency, this confirms that these rhythms can be differentiated on a cycle-by-cycle basis using their waveform shape. Additionally, we found that both alpha and mu rhythms increase in frequency and amplitude across development while becoming more nonsinusoidal. We further demonstrated that the two rhythms change differentially across age. Still, accounting for the significant changes in waveform features across development, we find no significant difference in waveform features for ADHD or ASD in comparison to typically developing participants, suggesting that no sensitive biomarkers can be constructed from these specified features of resting-state alpha-band rhythms.

## Materials and Methods

### Experimental design

We analyzed an openly available dataset from the Healthy Brain Network project (Alexander et al., 2017). The Healthy Brain Network project by the Child Mind Institute is an ongoing initiative aiming to generate a large-scale, transdiagnostic sample for biomarker discovery and for investigations of the neural substrates associated with commonly occurring disorder phenotypes. In the following, we give a brief summary of the experimental methods focusing on the resting-state EEG components of the data collection that were used in this study, with full details regarding all acquired data (including fMRI data) given in the original publication.

### Participants

Participants were recruited through advertisements that encouraged participation of families who had concerns about psychiatric symptoms in their child. These advertisements were distributed throughout the New York City metropolitan area. This recruitment strategy was developed with the main goal of the Healthy Brain Network–generating a large-scale, transdiagnostic sample–rather than the alternative of a fully representative epidemiologic design. Given this recruitment strategy, we have more participants with ADHD and/or ASD compared to those without a diagnosis than would be expected if recruitment was random. Participants completed several diagnostic assessments which were used to derive a consensus diagnosis by the lead clinicians, primarily based on the DSM-5-based Schedule for Affective Disorders and Schizophrenia - Children’s version (KSADS) psychiatric interview. As a part of the HBN protocols, participants completed a battery of 80+ forms of assessment across 12 hours and four testing sessions with a team of certified clinicians. As such, the diagnoses (or lack thereof) represent thorough and comprehensive testing for these disorders. Written consent was obtained from the participants’ legal guardians and written assent was obtained from the participants. Exclusion criteria for participation in the study were serious neurological disorders that prevented full participation in the complete study protocol (e.g. chronic epilepsy or low-functioning/non-verbal autism), as well as acute encephalopathy, known neurodegenerative disorder, existing formal diagnosis of schizophrenia, schizoaffective disorder or bipolar disorder, with a full list of exclusion criteria given in the original publication. Out of the 4245 participants listed in the first ten data releases, we were able to successfully download resting-state EEG data for 2969 participants, as resting-state EEG data acquisition was not conducted for all participants. To summarize the participant exclusion criteria (with description of the specific analysis steps following in subsequent paragraphs): we detected an alpha-band peak in at least one occipital or central channel in 2764 participants. 6 participants did not have an alpha peak in any of the 10 highest-SNR SSD components. 2273 participants had at least one source localized to alpha or mu regions, 1605 of these participants had sources that met the SNR and pattern distance thresholds (see subsection ***Classifying alpha rhythms as occipital alpha and sensorimotor mu*** subsection below) and were used for subsequent analysis of waveform features. Of those, 1247 were male and 358 were female. This sex distribution is consistent with the dominance of ADHD and ASD in the dataset, which have been reported to have greater prevalence in males than females, with 3:1 and 4:1 male/female ratio in children, respectively (Danielson et al., 2018; Baio, 2018). Bifurcating these 1605 participants by rhythm type detected, we isolated both alpha and mu rhythms from 238 participants, only an alpha rhythm from 1200 participants, and only a mu rhythm from 167 participants. The splits of these participants counts by diagnosis are enumerated in Table 1. The percent retention for each pipeline step was comparable across diagnoses.

**Table 1:** Participant counts by pipeline stage and diagnosis. Raw participant counts at each stage of the pipeline. The retention percentage relative to the number of participants listed online for that diagnosis.

| Condition | ADHD | ASD | No diagnosis given | Other diagnosis | Total |
| --- | --- | --- | --- | --- | --- |
| participant listed online | 1607 (100.0%) | 285 (100.0%) | 334 (100.0%) | 1759 (100.0%) | 4245 (100.0%) |
| EEG resting state available | 1224 (76.2%) | 201 (70.5%) | 263 (78.7%) | 1127 (64.1%) | 2969 (69.9%) |
| alpha-band peak detected, SSD run | 1143 (71.1%) | 191 (67.0%) | 241 (72.2%) | 1051 (59.7%) | 2764 (65.1%) |
| peak in SSD component detected | 1140 (70.9%) | 191 (67.0%) | 241 (72.2%) | 1048 (59.6%) | 2758 (65.0%) |
| source in alpha/mu regions, bycycle run | 956 (59.5%) | 161 (56.5%) | 194 (58.1%) | 851 (48.4%) | 2273 (53.5%) |
| Meet SNR and pattern distance thresholds | 704 (43.8%) | 117 (41.1%) | 139 (41.6%) | 573 (32.6%) | 1605 (37.8%) |

### EEG recordings

High-density EEG data were recorded throughout the session in a sound-shielded room using a 128-channel EEG geodesic hydrocel system by Electrical Geodesics Inc. The data was recorded at the sampling rate of 500 Hz and was bandpass filtered between 0.1 and 100 Hz. The recording reference was at the vertex of the head (electrode Cz), and impedances were kept below 40 kOhms throughout the recording. The participants underwent a series of tasks in the session. For this paper, we analyzed data from the resting-state paradigm, in which participants alternated between viewing a fixation cross with their eyes open for 20 seconds and closing their eyes for 40 seconds for a total of five minutes. Data analysis was performed with Python using MNE-Python, version 1.1.0 (Gramfort et al., 2013; Larson et al., 2022).

### Data preprocessing

We rejected channels that had values of zero for every time point (i.e., flat channels) and channels whose variance was greater than three standard deviations away from the channel’s mean value across time (i.e., had a high level of noise). We then used spherical spline interpolation for these rejected channels. We performed minimal preprocessing because SSD is able to separate components with high alpha SNR from the artifacts present in minimally processed EEG sensor data (Nikulin, Nolte, and Curio, 2011).

### Calculation of spectral parameterization

To calculate power spectra from the extracted time series, we used Welch’s method (window length: 4 seconds, 0% overlap, Hamming window). We then used the spectral parameterization toolbox (version 1.0.0) (Donoghue, Haller, et al., 2020) for the calculation of alpha oscillatory power and aperiodic exponents. The power spectrum is modeled as a combination of aperiodic and oscillatory components, which allows for isolation of alpha oscillatory power from aperiodic components. The power spectral density *P* (*f*) for each frequency *f* is expressed as:

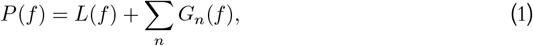

with the aperiodic contribution *L*(*f*) expressed as:

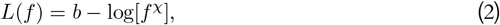

with a constant offset *b* and the aperiodic exponent *χ*. When the power spectrum is plotted on a log-log axis, the aperiodic exponent *χ* corresponds to the slope of a line. Each oscillatory contribution *G*_*n*_(*f*) is modeled as a Gaussian peak:

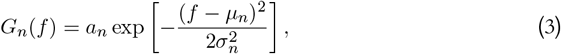

with *a*_*n*_ as the amplitude, *µ*_*n*_ as the center frequency and *σ*_*n*_ as the bandwidth of each component. The number of oscillatory components is determined from the data, with the option to set a maximum number of components as a parameter. The general model assumption here is that oscillatory and aperiodic processes are distinct and separable. Parameters for spectral parameterization were set as follows: peak width limits: (1, 12); maximum number of peaks: 5; minimum peak amplitude exceeding the aperiodic fit: 0.0; peak threshold: 2.0; and aperiodic mode: ‘fixed’.

Peak frequencies can vary across participants, and the success of spatial filters (described in the ***Calculation of spatial filters and patterns*** subsection) depend on filtering in a frequency range that includes the peak frequency of the rhythm one aims to isolate. Because of these constraints, it was important that we approximated each participant’s alpha and mu peak frequencies to find spatial filters that properly isolated these alpha-band rhythms. We first computed the spatial Laplacian to accentuate the local features in the data and reduce the effects of volume conduction; we were particularly concerned with reducing the effect of occipital alpha rhythms in sensorimotor channels so that we could estimate mu peak frequency. Next, we used spectral parameterization to find the peak frequency in the 6–13 Hz range for two occipital (E70 and E83, O1 and O2, respectively) and two sensorimotor channels (E36 and E104, C3 and C4, respectively) after computing the spatial Laplacian for these channels. This was done to get an approximate estimate of the peak frequencies for alpha and for mu rhythms in each participant to be used as a parameter in the spatial filter calculation described below. Of the 2969 participants for which we were able to successfully download their EEG data, 2764 had an alpha peak in at least one of the four selected channels. After alpha and mu components were identified, spectral parameterization was again applied to determine the peak frequency for each component to safeguard against variations in peak frequency. The aperiodic component was not used for any subsequent analyses, as the spatial filtering only optimizes for extracting oscillations and the effects of this procedure on the aperiodic exponent have not been characterized. Beyond this methodological constraint, developmental trajectories of the aperiodic exponent have been well-documented in previous studies (Wilkinson et al., 2024; Hill et al., 2022; Cellier et al., 2021), including with this developmental dataset (Caffarra et al., 2024; Tröndle et al., 2022).

### Calculation of spatial filters and patterns

For extracting rhythms in the alpha-band, we used spatio-spectral decomposition (SSD) (Nikulin, Nolte, and Curio, 2011). SSD maximizes spectral power in a frequency band of interest while minimizing spectral power in flanking frequency bands, thus enhancing the height of spectral peaks over the aperiodic component and exploiting the typical narrowband peak structure of neural rhythms (see Fig. 1C). Because alpha peak frequency can vary substantially across participants, we used the individual alpha peak frequency from spectral parameterization for each participant (*α*_freq_) as the center frequency for the SSD frequency band of interest. SSD models the measured time series **X** (a matrix with *t* samples and *k* electrodes) as a linear combination of signal **X**_**S**_ and noise **X**_**N**_:

**Figure 1:**
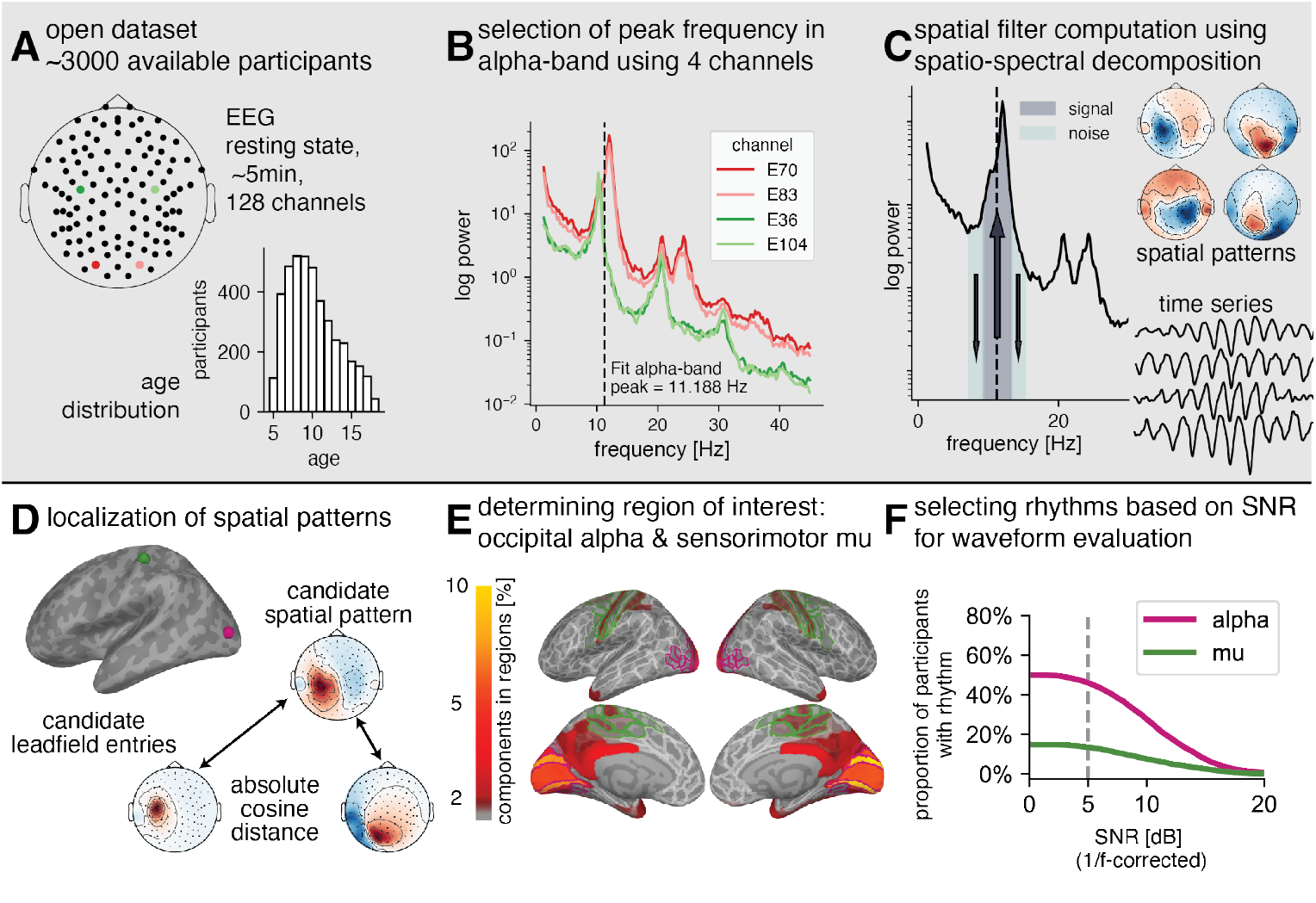
Analysis pipeline for extracting occipital alpha and sensorimotor mu rhythms with high SNR in a large developmental EEG dataset. (A) Sensor positions of 128-channel EEG (left) and the distribution of ages (bottom right) within the available dataset. (B) Power spectra for two occipital channels (red) and two sensorimotor channels (green) for one example participant in order to estimate the average indiviudal alpha-band peak frequency across the four channels (black dashed line). (C) The spatial filtering algorithm SSD extracts oscillatory sources with high alpha SNR by maximizing power within the defined frequency band of interest and minimizing power within defined neighboring frequency bands (left), with each source characterized by its spatial distribution (top right) and time series (bottom right). (D) To localize sources, the absolute cosine distance was computed for each SSD spatial pattern and lead field entry for each possible location. (E) Heat map of locations of the extracted alpha-band components superimposed on a 3D brain, for an SNR threshold of 5 dB and a pattern distance metric of 0.15. Components localized to an occipital region (outlined in pink) were classified as alpha components, while components localized to a sensorimotor region (outlined in green) were classified as mu components. (F) The percentage of participants who had a detectable rhythm exceeding a specified SNR threshold for alpha and mu rhythms, respectively.

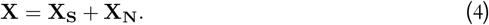

In our case, the signal **X**_**S**_ is the signal in the alpha band, while the noise **X**_**N**_ is the signal in the neighboring frequency bands. **X**_**S**_ was calculated by bandpass filtering (*α*_freq_ − 2, *α*_freq_ + 2). Following the procedure of Nikulin and colleagues (Nikulin, Nolte, and Curio, 2011), **X**_**N**_ was calculated by bandpass filtering (*α*_freq_ − 4, *α*_freq_ + 4) and then bandstop filtering (*α*_freq_ − 3, *α*_freq_ + 3). We can then calculate the signal covariance **C**_**S**_ and noise covariance **C**_**N**_ from the corresponding time series:

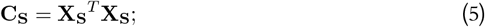

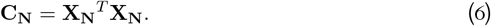

We aim to find spatial filters **W** that maximize the power of the projected signal *P*_*S*_ while minimizing the power of the projected noise *P*_*N*_ :

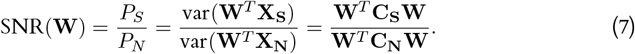

We can then transform this Rayleigh quotient into the following generalized eigenvalue problem:

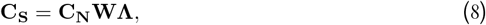

where **Λ** is the unity matrix with the corresponding eigenvalues on the diagonal. Each column of **W** is a spatial filter, and each participant will have as many spatial filters as electrodes (i.e., *k* spatial filters), with the spatial filters ordered by relative SNR in the frequency band of interest. Time series for each SSD component can be obtained from broadband activity across electrodes through matrix multiplication of the transposed spatial filters:

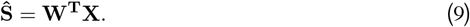

Crucially, because the SSD component time series is a linear transformation on the broadband activity rather than filtered activity, nonsinusoidal features of the waveforms, such as harmonics, are retained in the SSD component time series, allowing for fine-grained analysis of waveform shape. Finally, we can determine the spatial patterns **A** for each participant by inverting the matrix of spatial filters:

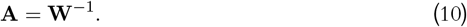

These spatial patterns can then be used to interpret the spatial origin of the extracted SSD components (Schaworonkow and Voytek, 2021a), and in our case, classify the SSD components as occipital alpha or sensorimotor mu (see below). The initial spatial patterns occasionally contain outlier channels that either have very large values or very different values from their neighboring channels within a spatial pattern that could corrupt source localization. Thus, we identified outlier channels programmatically for each participant by detecting channels that were three standard deviations away from the mean in the spatial pattern or the spatial derivative of the spatial pattern. Because these outlier channels are consistent across spatial patterns, we looked only at channels that were identified as outliers across at least three of the top 10 spatial patterns for each participant. We then recomputed SSD with each participant’s set of outlier channels removed. The SSD procedure generates three outputs for each component: a spatial filter, a spatially filtered time series, and a spatial pattern. The spatial pattern is used for the source localization described in the next section.

### Classifying alpha rhythms as occipital alpha and sensorimotor mu

To estimate source locations for each component, we computed the leadfield from a pediatric template brain and found the source that best accounts for that component’s spatial pattern. We used the atlas from Fonov and colleagues (Fonov et al., 2011), which averages participants MRIs in the age range of 4.5 to 18.5 years and used this as our template head and brain model. We then segmented the template head into the inner skull, outer skull, and outer skin surfaces and used these surfaces to create a 3-layer Boundary Element Method (BEM) model using FreeSurfer (Fischl, 2012). We aligned the FreeSurfer surfaces and BEM model to the template electrode positions. Next, we created a surface-based source space with a recursively subdivided octahedron. We computed the forward solution using the BEM model and surface-based source spaces, resulting in a spatial pattern for each dipole source. Finally, we computed the absolute cosine distance between each component’s spatial pattern and the spatial patterns generated by each dipole. The dipole with the lowest cosine distance was then defined as the source location for that component. Using the HCP-MMP1 parcellation (Glasser et al., 2016), components that were localized to the occipital cortex were classified as alpha components and those that were localized to somatosensory, motor, premotor, paracentral lobular, and midcingulate cortices were classified as mu components. 2216 participants had at least one component with a source classified as mu or alpha according to spatial location. To ensure that components were strong alpha sources, we calculated the aperiodic-adjusted SNR (spectral alpha peak height without the 1/f-contribution) for each component and excluded components with SNR ≤ 5 dB. We further defined each remaining component’s spatial pattern distance as the absolute cosine distance between that component’s spatial pattern and the spatial pattern generated by its source dipole. This spatial pattern distance quantifies how much the spatial pattern deviates from a noise-free leadfield pattern resulting from a dipole at the exact source location. To ensure that source localization was accurate, we excluded components with spatial pattern distance ≥ 0.15. Together then, these thresholds ensured that the components used for subsequent waveform shape analyses were strong alpha-band sources with well-estimated source locations. 1605 participants had at least one alpha or mu component that passed these SNR and spatial pattern distance thresholds. To simplify comparisons across participants, we only used each participant’s highest-SNR alpha component and highest-SNR mu component in further analyses.

### Calculation of waveform measures

For all of the alpha and mu components, we then calculated measures of waveform shape from the component time series using the bycycle package (version 1.0.0) (Cole and Voytek, 2019). This complementary approach to spectral analyses allows us to examine the nonsinusoidal, asymmetric features of the alpha and mu rhythms. First, we filtered the component time series in the broad frequency band from 1–45 Hz to remove slow drifts as well as high-frequency noise that would complicate extrema localization without deforming the shape of the alpha and mu rhythms. To extract the time points of zero crossings, the time series is then narrowband filtered at the component peak frequency identified earlier using spectral parameterization ± 2 Hz. Next, the peaks and troughs for each cycle are extracted by finding the absolute maxima and minima of the broadband signal between the zero-crossing time points. The rise and decay were defined as the phase from trough to peak and the phase from peak to trough, respectively. Because oscillations are infrequent and bursty (Cole and Voytek, 2019; Jones, 2016; Lundqvist et al., 2016), we next performed burst detection to determine which individual cycles represent alpha and mu bursts. Burst detection parameters were set as follows for all components: amplitude fraction threshold: 0.5 (cycles with amplitudes smaller than the median amplitude were discarded), amplitude consistency threshold: 0.5 (cycles where the difference in rise and decay voltage values was greater than the median difference were discarded), period consistency threshold: 0.5 (cycles where the difference between the cycle period and the period of neighboring cycles was greater than the median were discarded), monotonicity threshold: 0.5 (cycles where the fraction of instantaneous voltage changes between consecutive samples that are positive during rise and negative during the decay phase was smaller than 0.5 were discarded). These relatively stringent burst criteria have been used previously to investigate waveform shape of low-frequency bursts in infants (Schaworonkow and Voytek, 2021b). We determined the cycle frequency, rise-decay asymmetry, peak-trough asymmetry, and cycle amplitude from the times and voltages of the extracted critical points for each cycle as follows:

- frequency: 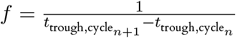
- rise-decay asymmetry: 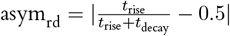
- peak-trough asymmetry: 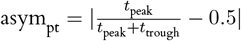
- amplitude: 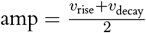

To account for any unequal voltage scaling across components from SSD, we normalized the cycle amplitude across each component by dividing the amplitude of cycles within bursts by the average amplitude of all cycles from non-bursts.

To separate the eyes open and eyes closed conditions, we extracted event markers for the opening and closing of eyes from the EEG signals. Because the modulation of alpha by closing the eyes can often take 1–2 seconds, we classified time periods two seconds after the closing of the eyes until two seconds before the subsequent opening of the eyes as eyes closed, and vice versa for eyes open. We then recomputed burst detection on the time series data for eyes closed and eyes open conditions separately because the relative amplitude criterion used therein could miss lower amplitude alpha bursts in the eyes open condition.

### Statistical analysis

Statistical analyses were carried out using the pingouin Python package (version 0.5.3) (Vallat, 2018). We used *t*-tests to determine if there were significant differences in waveform features between alpha and mu (Fig. 2). We evaluated the effect sizes of these contrasts using Cohen’s *d*. In order to determine whether the waveform features change across development for alpha and mu, we first calculated correlation coefficients across age for each rhythm type (Fig. 3A). To assess whether the waveform features change differently by rhythm type, we used permutation statistics and linear models to bootstrap the slope and offset for each waveform feature of each rhythm type. Specifically, we resampled feature values with replacement and fit a linear model for each rhythm type in which each feature is a function of age: feature = *β*_1_ · age + *β*_0_, giving us bootstrapped distributions of slope (*β*_1_) and offset (*β*_0_) for each waveform feature and rhythm type (Fig. 3B-C). We then calculated *p*-values for differences in the *β* coefficients between rhythm types by comparing the percentage of bootstraps where the difference of the *β* coefficient of alpha and mu was either above or below zero to 50%. Finally, we evaluated whether waveform shape features differ across clinical diagnoses (Fig. 4), independent of the observed changes in these waveform features with age, by performing an ANCOVA and using age as a covariate. We performed post-hoc *t*-tests on the residuals obtained from the linear regression of age with each waveform feature. This analysis aimed to verify pairwise non-significance in line with the results of the ANCOVA. To determine whether the diagnoses data constitute strong evidence for the null hypothesis, we calculated Bayes Factors (BFs; Jeffreys, 1998; Wetzels et al., 2011) using a JZS (Jeffreys-Zellner-Siow) prior (Rouder et al., 2009). According to Jeffreys’ scheme,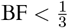 and 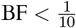 indicate substantial and strong evidence for the null hypothesis, respectively,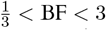 indicates insufficient evidence, and BF *>* 3 and BF *>* 10 indicate substantial and strong evidence for the alternative hypothesis, respectively (Jeffreys, 1998; Jarosz and Wiley, 2014). All reported *p*-values were corrected for multiple comparisons using the Bonferroni correction, except in the case of relating waveform features to diagnoses, where we used the Benjamini-Hochberg procedure (Benjamini and Hochberg, 1995) to decrease the chance of Type II errors (i.e., falsely claim that there are no significant differences in waveform features across diagnoses when there are).

**Figure 2:**
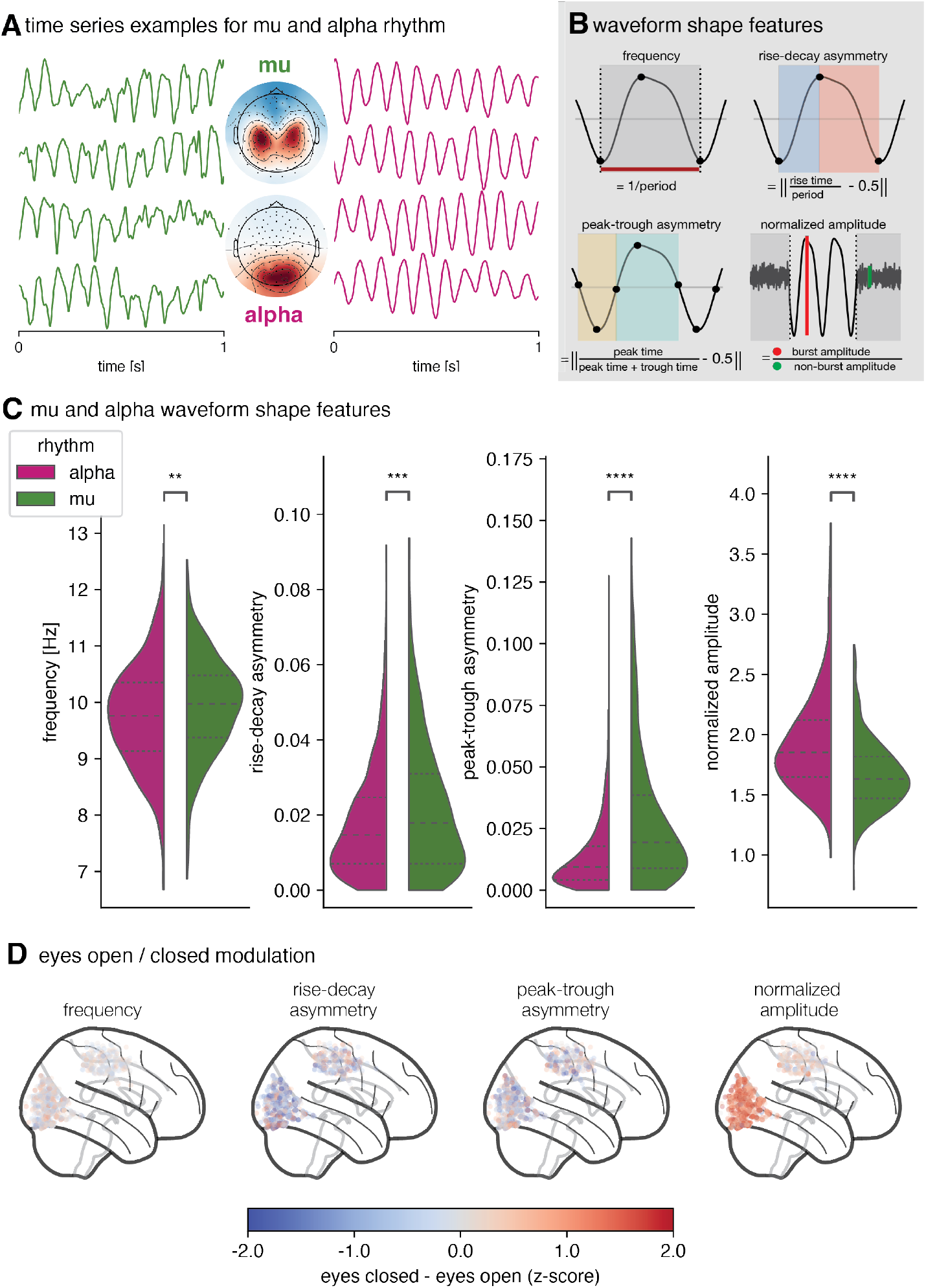
Waveform features differ between alpha and mu rhythms. (A) Example time series for alpha (right) and mu (left) components from two participants. The topoplot inset shows the average topography for each rhythm type across all included participants (top: mu, bottom: alpha. (B) Schematic illustrations of assessed waveform features: frequency, rise-decay asymmetry, peak-trough asymmetry, and normalized amplitude. (C) Estimated distribution of mean waveform features across time for extracted alpha rhythms (pink) and mu rhythms (green) across participants (N=1605). (D) Locations of alpha (posterior cluster) and mu (central cluster) sources showing modulation across the eyes closed and open conditions for the four examined features (eyes closed minus eyes open). The c1o3lor map was adjusted, so that the color limits correspond to z-scored differences of 2 and −2.

**Figure 3:**
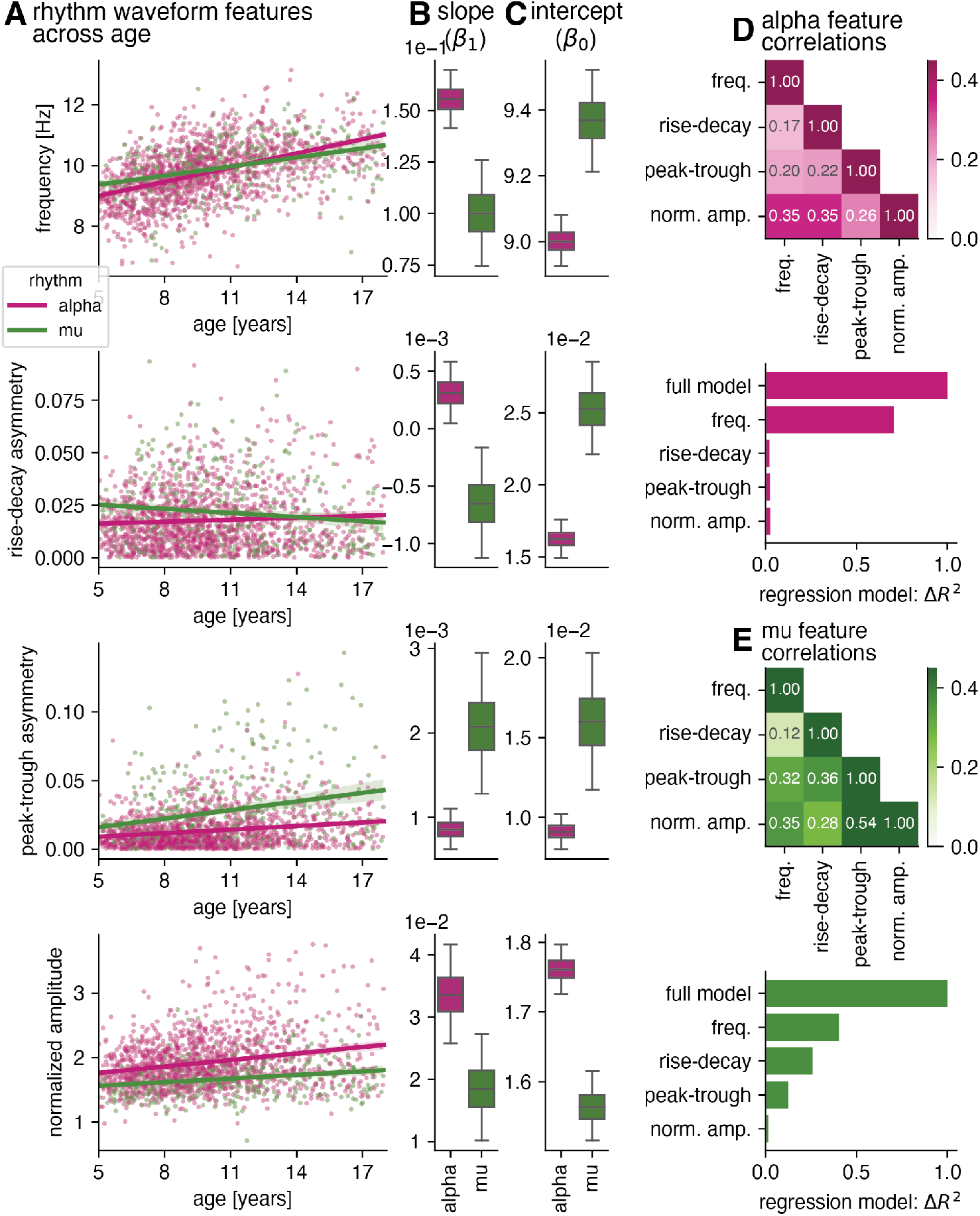
Alpha and mu rhythm waveform features show differential changes across development, analysis collapsed across eyes open and eyes closed conditions. (A) Alpha (pink) and mu (green) waveform features (frequency, rise-decay and peak-trough asymmetry, normalized amplitude) by age across participants. The scatterplots for both rhythm types individually can also be seen in Fig. C1, providing improved distinction. (B) Slope of linear model for alpha and mu rhythm. Boxes denote interquartile range, and whiskers denote 95% confidence interval of bootstrapped feature values (10,000 iterations). (C) Intercept of linear model for alpha and mu rhythm. Boxes denote interquartile range, and whiskers denote 95% confidence interval of bootstrapped feature values (10,000 iterations). (D) alpha rhythm: (top) Spearman correlation between mean waveform features across participants; (bottom) unique contribution of individual waveform features to age in a multivariate linear regression model, as estimated by variance partitioning. (E) same as (D) but for the mu rhythm.

## Results

### Methodological approach for establishing the presence of alpha and mu rhythms

Our processing pipeline aimed for a computationally efficient, noise-resistant investigation of EEG alpha-band rhythms without any manual data cleaning in an EEG dataset of nearly 3000 minors aged 5–18 (mean age: 10.1 ± 3.2 years), consisting of approximately 5 minutes of resting-state activity in 128 channels (Fig. 1A). Our focus was specifically occipital alpha rhythms and sensorimotor mu rhythms. First, we identified the peak frequency in a broad, 6–13 Hz range in two occipital and two sensorimotor channels for each participant using the spectral parameterization method (Donoghue, Haller, et al., 2020) (Fig. 1B). Alpha-peak detection and spectral parameterization model goodness-of-fit metrics were consistently high across the ages investigated (Fig. A1 and Fig. A2). We then used each individual’s average alpha-band peak frequency across the four channels to seed a spatial filtering algorithm called spatio-spectral decomposition (SSD) that extracts oscillatory sources with high alpha-band signal-to-noise ratio (SNR). Each extracted source consisted of a time series and associated spatial topography (Fig. 1C). These SSD sources attenuate noise in higher frequencies that is apparent in the sensor-space EEG (Fig. A3). To maximize accuracy of source localization for these rhythms (described below), we performed outlier detection on the initial spatial patterns and recomputed SSD with any detected outlier channels removed. Using a template magnetic resonance image developed specifically for a developmental population (Fonov et al., 2011), we used a template matching approach to estimate the source location that best accounts for the spatial pattern of the strongest components with high alpha-band SNR for each participant (Fig. 1D). The spatial patterns were compared to the leadfield of 32,772 source locations using a cosine-based distance metric. The majority of sources were localized to occipital and sensorimotor cortices after imposing alpha-band SNR and acceptable pattern distance thresholds (Fig. 1E). We classified the SSD components with SNR > 5 dB located in the occipital cortex as alpha rhythms (Fig. 1E- F, pink) and those located in sensorimotor cortices as mu rhythms (Fig. 1E-F, green). Our extraction procedure resulted in sensorimotor mu rhythms for 13.6% (405/2969) of participants and visual alpha rhythms for 48.4% (1438/2969) of participants (Fig. 1F). This percentage for detected mu rhythms is comparable to that achieved by previous studies (Aird and Y Gastaut, 1959; Koshino and Niedermeyer, 1975; Frauscher et al., 2018), while the percentage of detected alpha rhythms reflects the fact that we intentionally excluded parietal alpha rhythms, which are prominent in resting-state, sensor-space EEG (Sokoliuk et al., 2019). The time series for these selected high-SNR alpha and mu rhythms were then used for all subsequent waveform shape analyses.

### Alpha and mu rhythms differ in their oscillatory waveform features

After extracting alpha and mu rhythm time series (example seen in Fig. 2A), we computed waveform features on a cycle-by-cycle basis, using the same approach as (Cole and Voytek, 2019). Each time series was divided into cycles by identifying zero crossings and extrema (peaks and troughs) for each cycle of the alpha-band rhythm. A cycle was classified as a burst after passing burst detection criteria (see **Methods**). We quantified four waveform features: average frequency, rise-decay asymmetry, peak-trough asymmetry, and amplitude (see Fig. 2B). The oscillation frequency was computed as the inverse of the cycle period. We assessed the asymmetry of the time series with two different metrics, using the rise-decay asymmetry as well as the peak-trough asymmetry; the former measures the asymmetry in the amount of time spent in the oscillation cycle rise versus decay, whereas the latter measures the amount of time spent in the oscillation peak compared to the trough, approximating the “sharpness” of the cycle extrema. Normalized amplitude was computed as the ratio between the amplitude of the burst vs. non-burst cycles. This measure strongly correlates with the 1/f-corrected spectral peak alpha-band amplitude (*r* = 0.844), but has the benefit that it can be calculated from the time series itself.

Because both types of rhythms could not be extracted for every participant, we assessed all measures both across and within participants. Across-participants analyses maximized the number of participants included in the analysis to ensure observed differences in alpha and mu rhythms were representative of the participants as a whole, while within-participant analyses ensured any differences seen across participants reflected differences in rhythms rather than any other characteristic of participants that varied across rhythm types (e.g., age). Comparisons of alpha and mu waveform features between participants are shown in Fig. 2C: alpha peak frequency was lower than mu peak frequency (*t* = − 3.71, *d* = − 0.200, *p* = 0.00182). Additionally, mu rhythms were more asymmetrical than alpha rhythms in both measures of asymmetry: rise-decay asymmetry (*t* = − 4.08, *d* = − 0.257, *p* = 4.09 · 10^−4^) and peak-trough asymmetry (*t* = − 10.8, *d* = − 0.870, *p* = 1.39 · 10^−23^). Normalized amplitude was higher for alpha rhythms (*t* = 14.6, *d* = 0.668, *p* = 1.56 · 10^−42^), reflecting burst cycles of higher magnitude. Within participants, waveform features showed the same alpha/mu differences as between participants, such that alpha rhythms showed lower peak frequency, less asymmetry, and were of higher amplitude than mu rhythms. Effect sizes within participants were also of similar magnitude to those across participants (Table B1). This shows that effects seen across the population are also generally reflected within individuals. This consistency also indicates that the observed differences are not driven by participants for which only one type of rhythm could be detected, nor by differences in sample size between alpha and mu rhythms. As the dataset included periods of eyes open/eyes closed modulation, we were interested whether our pipeline would reflect commonly known amplitude modulation of alpha during eyes closed periods, while maintaining a stable amplitude for mu. Fig. 2D shows eyes open/eyes closed modulation for different features across the brain. Consistent with previous studies, alpha amplitude was significantly decreased from eyes closed to eyes open (*t* = 48.2, *d* = 1.39, *p* = 3.02 · 10^−296^), with a much larger effect size than the difference in mu amplitude between eyes closed and eyes open (*t* = 4.53, *d* = 0.124, *p* = 6.50 · 10^−5^). This result bolsters the notion that our pipeline properly isolates alpha and mu rhythms. Frequency was not significantly different between eyes closed and eyes open for alpha (frequency: *t* = 2.42, *d* = 0.0328, *p* = 0.124) nor mu (*t* = − 0.711, *d* = − 0.0125, *p* = 1). Waveforms were significantly more asymmetrical during eyes open than during eyes closed for both alpha (rise-decay asymmetry: *t* = − 16.2, *d* = − 0.506, *p* = 2.28 · 10^−53^; peak-trough asymmetry: *t* = − 7.93, *d* = − 0.267, *p* = 3.49 · 10^−14^) and for mu (rise-decay asymmetry: *t* = − 4.95, *d* = − 0.252, *p* = 9.20 · 10^−6^; peak-trough asymmetry: *t* = −5.29, *d* = −0.182, *p* = 1.67 · 10^−6^). With these increases in waveform asymmetry being relatively small in comparison to the decrease in alpha amplitude, the strongest effect of opening the eyes remains the decrease in alpha amplitude that is consistent with previous results. As the mu amplitude remained stable across eyes open/eyes closed periods and the eyes closed alpha cycles contributed the majority of alpha cycles, we collapsed across eyes open and closed periods for further analyses.

In summary, the results indicate that our analysis pipeline was able to extract differential alpha and mu rhythm waveform characteristics effectively without any manual data cleaning in a large dataset. Moreover, these results align with qualitative descriptions in previous literature regarding oscillatory frequency and amplitude and provide a novel, quantitative characterization of waveform shape, distinctly for both rhythm types.

### Waveform features differ strongly across development

After evaluation of waveform shape features for each rhythm type, we next investigated how these waveform shape features change across development. The results are shown in Fig. 3A. First, we reproduced the classic increase of frequency across development for both alpha (*r* = 0.518, *p <* 10^−80^) and mu (*r* = 0.421, *p* = 3.47 · 10^−18^). Second, mu rise-decay asymmetry decreases significantly across development (*r* = − 0.125, *p* = 0.047), while alpha does not show a significant increase (*r* = 0.017, *p* = 1.0). For peak-trough asymmetry, both rhythm types display increasing nonsinusoidality with higher age (alpha: *r* = 0.167, *p* = 6.87 · 10^−10^; mu: *r* = 0.249, *p* = 1.56 · 10^−6^). Additionally, we showed that both alpha (*r* = 0.217, *p* = 4.08 · 10^−16^) and mu (*r* = 0.244, *p* = 2.78 · 10^−6^) increase in burst amplitude relative to non-burst cycle amplitude across development.

To compare the relationship between waveform features and age across the two rhythm types in a more fine-grained way, we estimated linear models using resampling and permutation statistics. We formulate a linear model for each rhythm type in which each feature is a function of age: feature = *β*_1_ · age + *β*_0_. To determine whether there are distinct slopes and offsets for mu and alpha rhythms in their relationship with age, we performed resampling with replacement of feature values and fit linear models to the resampled feature values across age for both rhythm types to calculate *p*-values (results are reported as 95% confidence intervals of the difference between alpha and mu parameters, in square brackets). Our results show that the two rhythm types change differentially across age (see Fig. 3B for slope and Fig. 3C for intercept). The slope differences indicate that changes in rhythm characteristics across development affect one rhythm more than the other. Alpha changes across development were more pronounced than those for mu in frequency ([0.0259, 0.0852], *p* = 0.002), rise-decay asymmetry ([0.0004, 0.0015], *p* = 0.0006), and amplitude ([0.0032, 0.027], *p* = 0.0126), while changes in peak-trough asymmetry were more pronounced for mu than alpha ([−0.0021, −0.0004], *p* = 0.0048). The difference in intercept indicates that the changes in rhythm characteristics are already evident at the beginning of the age range in this study (minimum age 5). Frequency ([−0.5412, −0.192], *p* = 0.0002) and both asymmetry measures (rise-decay: [−0.0126, −0.0056], *p <* 0.0001; peak-trough: [−0.0113, −0.0024], *p* = 0.004) were higher for mu at age 5, while amplitude ([0.1347, 0.2596], *p <* 0.0001) was higher for alpha at age 5.

To investigate whether the changes in waveform features with age are nonlinear, we fit generalized additive models (GAMs) to these data. We qualitatively observe a close correspondence within the investigated age range (5–18 years) to the linear model (Fig. C1), validating the use of linear models. Furthermore, as the data contained both eyes closed and eyes open data segments, we also performed this analysis separately for each type of segment. As shown in Fig. C2, the main results when concatenating across eyes open/closed primarily resemble the eyes closed condition. This is unsurprising, given that there are more data segments with eyes closed than eyes open (180 seconds vs. 120 seconds). That said, the changes across age for eyes open were qualitatively similar to the eyes closed condition for frequency and peak-trough asymmetry, with increasing waveform feature values throughout development. The differences between alpha and mu for rise-decay asymmetry were entirely driven by the closed condition, as there were no significant differences in slope or intercept between alpha and mu with eyes open. The normalized amplitude measure did not show a difference between eyes closed and eyes open conditions for mu, but the slope and intercept were higher for eyes closed than eyes open for alpha. Finally, to account for potential differences in age across rhythm types, we fit linear models with bootstrapping to only the participants with both alpha and mu rhythms (Fig. C3). The linear model parameters are comparable to those for all participants, validating that the differences between alpha and mu waveform features across age are not due to the age distribution across the rhythm types.

The individual features are correlated to a varying degree (Fig. 3D). Therefore, to investigate if the individual waveform features capture unique information in the light of their correlations, we performed a unique contribution analysis. For this, we estimate a multivariate model predicting age from the four waveform features, calculating 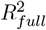 for the full model. The full model had an 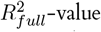 of 0.257 for the alpha rhythm and 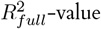 of 0.213 for the mu rhythm. Next, we removed one feature and estimated a linear model for age using the remaining three waveform features to obtain a 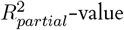. This was used to calculate the unique relative contribution of the remaining individual waveform feature: 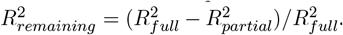. Relative contributions are shown in Fig. 3E: while for the alpha rhythm a large amount of information related to age is captured uniquely by the frequency alone, without waveform asymmetries being informative about age, for the mu rhythm a substantial amount of information related to age is captured uniquely by waveform asymmetries. For both rhythm types, normalized amplitude is not uniquely informative about age. Overall, this finding highlights the differing roles of specific waveform features in changes across age for both rhythm types, with the remaining predictive power arising from the shared information between waveform features.

In this way, our procedure was able to quantify differences in waveform features separately for mu and alpha, with each rhythm type displaying unique characteristics, albeit with high individual variance. Changes in waveform features across development for mu were invariant to eyes open/closed; while there were differences across eyes open and eyes closed for alpha, these differences reflect classical eyes open/closed effects. In the following section, we examine whether the individual variance can be linked to several types of clinical diagnoses.

### Resting-state rhythm features across diagnoses

Given the easy applicability of resting-state EEG in a developmental population, there is substantial interest in deriving sensitive biomarkers from resting-state EEG. Therefore, we compared the above waveform shape features across clinical diagnoses to see if alpha rhythm (Fig. 4A) or mu rhythm (Fig. 4B) features could be related to ADHD or ASD diagnoses. An analysis of covariance (ANCOVA) was conducted to examine the influence of diagnosis (with three levels: ADHD, ASD, and no diagnosis) on each of the four waveform features while accounting for differences in the covariate age across diagnostic groups. The main factor of diagnosis was not significant for any of the waveform features for both alpha (frequency: *F* = 2.649, *p* = 0.570, rise-decay asymmetry: *F* = 0.828, *p* = 0.583, peak-trough asymmetry: *F* = 0.886, *p* = 0.583, and normalized amplitude: *F* = 1.213, *p* = 0.583) and mu rhythms (frequency: *F* = 0.034, *p* = 0.967, rise-decay asymmetry: *F* = 0.863, *p* = 0.583, peak-trough asymmetry: *F* = 1.899, *p* = 0.583, and normalized amplitude: *F* = 0.498, *p* = 0.696). These null results were further supported by pairwise post-hoc *t*-tests that controlled for age. Furthermore, supplementary analyses showed that there were no significant differences between diagnoses when the data were split into the eyes open (Fig. D1) and eyes closed conditions (Fig. D2). Thus, taking age into account, there were no significant differences in all four waveform shape features we investigated between participants with ADHD and ASD compared to typically developing participants for either alpha-band rhythm. Given the non-significant findings for each of the waveform shape features across the diagnoses, we calculated Bayes factors (BFs) to examine statistical evidence for the null hypothesis that there are no difference in waveform shape features for alpha and mu rhythms in ADHD and ASD (Jeffreys, 1998). All 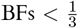, indicating strong evidence for the null hypothesis, except for frequency in ADHD vs. typically developing (BF = 0.835) and normalized amplitude in ASD vs. typically developing (BF = 0.440). Regarding developmental changes, there were no differences between diagnostic groups and participants who received no diagnosis in the regression coefficients relating age and waveform features for alpha (Fig. D3) nor mu (Fig. D4) rhythms.

**Figure 4:**
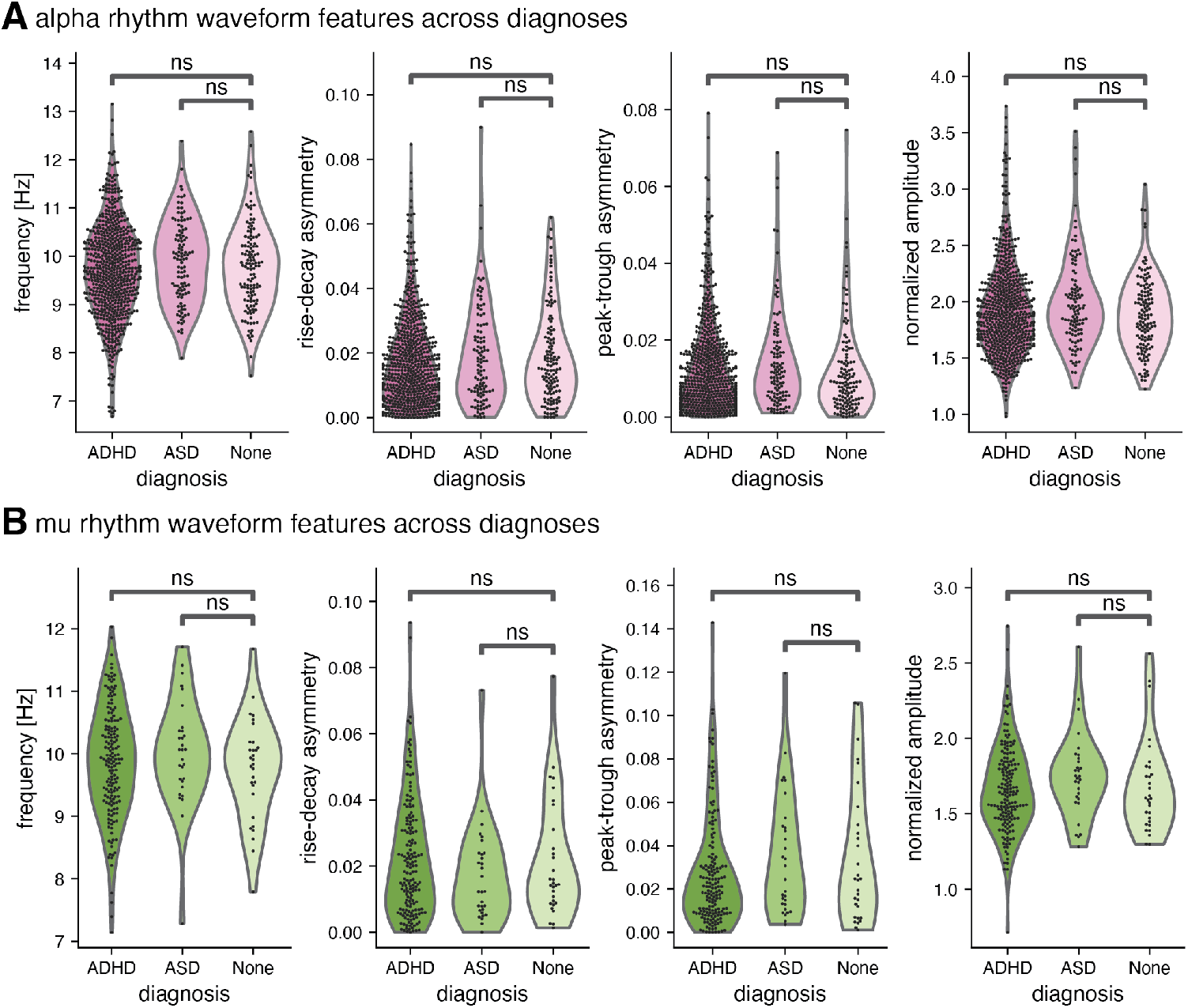
Resting-state alpha and mu waveform features across ADHD and ASD diagnoses compared to typically developing participants. (A) alpha rhythm components: mean waveform features (frequency, rise-decay and peak-trough asymmetry, normalized amplitude) in participants who received an ADHD and ASD diagnosis, and for participants who did not receive a diagnosis (ADHD: N=639, ASD: N=129, no diagnosis N=109). (B) mu rhythm components: mean waveform features in participants who received an ADHD and ASD diagnosis, and for participants without a diagnosis (ADHD: N=167, ASD: N=30, no diagnosis N=30). No statistically significant difference between the two disorder groups and typically developing participants was found for any waveform feature and rhythm type.

Participants classified as having a ADHD or ASD diagnosis exhibit high heterogeneity, showing a wide range of symptoms, cognitive abilities, and comorbidities. Given this diversity, we investigated whether the variability in clinical presentation was related to differences in waveform features, using additional available information within the dataset. For the ADHD group, we compared the Hyperactive/Impulsive Type, the Inattentive Type, and the Combined Type, as given per clinician diagnosis, and the participants without a diagnosis using an individual ANCOVA per feature with age as covariate, finding no significant effects for diagnosis subtype as a factor, see Table D1. Additionally, we assessed the relationship between Conners 3 questionnaire scores, an ADHD-specific assessment, and waveform features within the ADHD group and across all participants with available scores. No statistically significant associations were found across any of the waveform features examined for each rhythm type, see Table D2 and Table D3. In the ASD group, we assessed the relationship between Autism Spectrum Screening Questionnaire (ASSQ) scores and waveform features, using Spearman correlation, with and without age as a covariate, finding no significant correlation (Table D4). This was also the case when using the ASSQ scores for all participants for which these were available, and relating them to waveform features (Table D5). In terms of comorbidities, a notable subset of participants had secondary diagnoses in addition to their primary diagnosis. Specifically, the number of participants with a primary diagnosis of ASD and a secondary diagnosis of ADHD or vice versa, was 115 out of 748 participants with a ADHD or ASD diagnosis (15.4%) for alpha and 30 out of 197 (15.2%) for mu. To investigate whether this subgroup exhibits distinct characteristics, we categorized the participants into a separate group, and conducted an ANCOVA with age as covariate, mirroring the main analysis in Fig. 4. The results indicated no significant effect of diagnosis (see Table D6, Fig. D5, and Fig. D6), suggesting that the presence of comorbidities did not substantially alter the waveform features analyzed. To evaluate the influence of cognitive abilities, we examined available Wechsler Intelligence Scale for Children (WISC-V) scores of all possible participants for which the WISC-V was completed (N=958). The ANCOVA relating different WISC-V subscores and diagnoses was significant for the main factor of diagnosis (Table D7 and Fig. D7), and so we estimated a linear model for waveform feature relationships that includes WISC-V score as an additional factor in addition to age and diagnosis. This factor was not significant, see model summary in Table D8. In summary, there was no significant difference for all waveform features across diagnoses, even when controlling for a variety of different variables related to the heterogeneity of the population.

## Discussion

With these analyses, we have established a data analysis pipeline to extract distinct oscillatory rhythms that overlap in frequency in resting-state EEG. The data-driven spatial filtering approach leverages the differential spatiospectral distribution of occipital alpha and sensorimotor mu to identify independent rhythms, as there could be several underlying generators that are differentially altered (Dumas et al., 2014). This approach effectively identifies components that display the narrowband peak structure of neural alpha-band rhythms with significant alpha power exceeding 1/f-activity, in the presence of noise, reducing the need for extensive preprocessing. This approach also reduces the chance of conflation of oscillatory and 1/f-activity in this context (Cellier et al., 2021). We used these spatial filters to linearly transform the broadband activity across electrodes into component time series, enabling the quantification of waveform shape across rhythms, diagnoses, and development.

This analysis demonstrated that both occipital alpha and sensorimotor mu have distinct, non-sinusoidal waveforms. We replicated long-standing findings that occipital alpha has lower frequency and higher amplitude than sensorimotor mu at a large scale (up to 1605 participants). Through our time series analysis of waveform shape, we were able to quantify the observed qualitative differences between the waveform shape of occipital alpha and sensorimotor mu. Moreover, the waveform asymmetry differences we found were much stronger than the well-established differences in frequency and amplitude, suggesting that the principal difference between these rhythms lies in the distinct asymmetries of each rhythm’s stereotyped waveform. Both these rhythms show substantial changes in waveform shape across development, increasing in frequency and amplitude and becoming more nonsinusoidal. Differences in waveform features are already present at age five, and the rate at which these waveform features change across development is significantly different between alpha and mu. Despite the significant differences between the waveforms of alpha and mu, neither rhythm showed any statistically significant difference in waveform shape across the examined diagnoses.

### Implications

Foremost, our quantitative analyses show that a mu rhythm can be identified only in a fraction of participants, matching more qualitative descriptions of previous literature. This underscores the critical need to verify the existence of an oscillation (Donoghue, Schaworonkow, and Voytek, 2021) before oscillatory measures are subjected to further analysis to increase validity and reliability of results. If an oscillation is present only in a fraction of participants, there may be robust group-level effects, but it may be challenging to use this measure as a biomarker, given its lack of presence in individuals. As the mu rhythm is often investigated in a developmental context, this pertains to a number of developmental studies. While our results demonstrate the importance of verifying the existence of the mu rhythm in developmental studies, this consideration extends beyond studies of the mu rhythm across development and is of importance for all studies investigating changes in brain rhythms, given the variability in how they manifest within and across participants. Multiple different modeling approaches have been employed to understand the generation of alpha-band rhythms. Quantifying the waveform shape of the distinct rhythm types that can be measured in the human EEG will aid in constraining generative models of alpha-band rhythms, for which the underlying generators are still insufficiently understood (Halgren et al., 2019). Neural mass models have been used to represent the average membrane potential and firing rate of large numbers of neurons (Lopes Da Silva et al., 1976), generating rhythmic activity similar to biologically observed alpha rhythms (Jansen and Rit, 1995), and making it possible to connect features derived from data, such as peak frequency and waveform shape to physiologically interpretable model parameters, such as maximum postsynaptic potential amplitude, firing rates, or connection strengths between populations. To our knowledge, generative biophysical models for different types of alpha rhythms that take waveform shape into account have not yet been developed. Our findings show that both alpha and mu rhythms have stereotyped, nonsinusoidal waveforms that become more asymmetrical across development. Such robust descriptive statistics will aid in the constraining of future generative biophysical models for these rhythms, which should account for these differences in waveform shape, both across rhythms and across development.

While important for understanding electrophysiological data at the level of local field potentials and EEG, these models fail to explain rhythm genesis at the level of synapses and individual neurons, which constitutes a promising target for interventions that seek to ameliorate cognitive deficits associated with these rhythms. As a first step, we have provided a conceptual model of waveform shape and the relationship between spike synchrony and waveform asymmetry of local field potentials in recent work (García-Rosales, Schaworonkow, and Hechavarria, 2024). In this work, a relationship to measures derived from spiking activity (spike synchrony across units, spike-LFP phase relationships, and pairwise phase consistency) have been shown to relate to the waveform asymmetry when examined across regions. There has been work investigating oscillations across development in animal models (Bitzenhofer, Pöpplau, and Hanganu-Opatz, 2020; Caveness, 1962), but without investigation of waveform shape. Future animal studies could investigate regionally specific oscillations across development, focusing on two key aspects: first, whether oscillatory waveforms change during development, and second, whether these changes are related to the aforementioned spike activity measures. Descriptive studies across species and developmental stages will enable us to draw parallels between models of rhythmogenesis and establish correspondences between regionally specific rhythm types in humans and animals. Our findings add to an already-substantial amount of studies showing that developmental changes in oscillations are dramatic (Freschl et al., 2022), despite significant inter-individual variability in the precise timing of such changes. The structural and physiological changes that underlie these pronounced changes across development remain unknown. Understanding these structural and physiological underpinnings of alpha-band rhythms will likely be crucial for future interventions that aim to mitigate the functional deficits that have been correlated with alpha-band power differences. The developmental trajectories of structural brain characteristics, such as gray matter volume, white matter volume, and cortical thickness, have been well-documented (Bethlehem et al., 2022). In future studies, these developmental trajectories of structural brain features could be related to the development trajectories of alpha-band waveform features to establish candidate structural contributors to changes in alpha-band waveform that can be tested more rigorously. Furthermore, these changes in resting-state alpha-band waveform features may reflect changes in cognitive abilities across development, given the implication of alpha rhythms in working memory (Foster et al., 2016; Jokisch and Jensen, 2007; Ede, 2018) and attention (Klimesch, 2012; Snyder and Foxe, 2010) and the improvements across development in working memory (Gathercole et al., 2004; Kwon, Reiss, and Menon, 2002) and attention (Luna et al., 2004; Rueda, Posner, and Rothbart, 2005).

Contrary to work implicating beta waveform shape in Parkinson’s disease (Cole, van der Meij, et al., 2017), we found no significant differences in alpha-band rhythm features for either of two prominent neurodevelopmental disorders. This is consistent with recent work from Dede and colleagues (Dede et al., 2023) that failed to find a single biomarker for ASD after extracting a high number of measures from resting-state EEG (in sensor space, including band-power and frequency measures) in a dataset of 776 individuals. There is large inter-individual variation in alpha-band rhythm features, but high internal consistency (Lopez et al., 2023), which hints at the fact that usage of longitudinal data (Gabard-Durnam et al., 2019) may be a promising approach to capture diagnosis-related information sufficiently. There is well-documented heterogeneity within both ASD (Lord et al., 2018; Happé and Ronald, 2008) and ADHD (Nigg, Tannock, and Rohde, 2010; Sonuga-Barke and Halperin, 2010) populations. Given this heterogeneity, we conducted several control analyses, taking into account clinical subtypes, symptom severity, comorbidities, and cognitive abilities. None of these control analyses resulted in a significant difference in waveform features. Our analysis suggests that it is challenging to use single-session resting-state oscillations for the construction of predictive biomarkers in the case of ADHD and autism, but does not rule out that a more specific task-operationalization of deficits may be used to uncover disorder-specific physiology, e.g. in specialized social paradigms for autism or attentional paradigms for ADHD (Vollebregt et al., 2016; Ter Huurne et al., 2017; Oberman et al., 2005). In summary, the need to go beyond resting-state analyses in this context is highlighted here, in order to yield robust and reproducible results, which is a challenge in this context (Lefebvre et al., 2018).

### Limitations

There are two notable limitations regarding the recruitment strategy and exclusion criteria of the original paper. First, because of the recruitment strategy, those not given a diagnosis after going through the full Healthy Brain Network protocol may still present overlapping symptoms to those diagnosed with ADHD or ASD without meeting the diagnostic thresholds for either disorder. We addressed this concern through examination of waveform features across ADHD subtypes, scores in an ADHD-specific assessment (Conners 3 questionnaire), and scores in an ASD-specific assessment (ASSQ). While these analyses mirrored the main findings of no significant differences in waveform features across diagnoses, it remains possible that overlapping symptoms of those given no diagnosis reduce the ability to observe statistical differences in alpha and mu waveform features. Second, since the exclusion criteria included severe cognitive impairment, our findings do not pertain to children with intellectual disability. This is particularly notable in the case of autism, where 37.9% of school age children with autism have intellectual disability (Maenner, 2023). As such, the ASD participants do not reflect the full spectrum of autistic individuals.

There are two notable limitations regarding the recruitment strategy and exclusion criteria of the original paper. First, because of the recruitment strategy, those not given a diagnosis after going through the full HBN protocol may still present overlapping symptoms to those diagnosed with ADHD or ASD without meeting the diagnostic thresholds for either disorder. We addressed this concern through examination of waveform features across ADHD subtypes, scores in an ADHD-specific assessment (Conners 3 questionnaire), and scores in an ASD-specific assessment (ASSQ). While these analyses mirrored the main findings of no significant differences in waveform features across diagnoses, it remains possible that overlapping symptoms of those given no diagnosis reduce the ability to observe statistical differences in alpha and mu waveform features. Second, since the exclusion criteria included severe cognitive impairment, our findings do not pertain to children with intellectual disability. This is particularly notable in the case of autism, where 37.9% of school age children with autism have intellectual disability (Maenner, 2023). As such, the ASD participants do not reflect the full spectrum of autistic individuals.

Given the heterogeneous data quality of a developmental EEG dataset, we made several design choices in our processing pipeline. First, we focused on high-SNR alpha-band rhythms, discarding any rhythms with SNR < 5 dB. The size of the dataset allowed us to be conservative in our rhythm classification while maintaining high statistical power for comparisons across the two rhythm types. Such a stringent SNR criterion would severely limit the number of SSD components one could extract for other rhythms with lower SNR (e.g., beta rhythms) and might prove unnecessarily stringent for datasets that are inherently less noisy or have been more thoroughly preprocessed. Second, we discarded parietal alpha-band rhythms in this study. Occipital alpha and sensorimotor mu have well-established roles in modulation of sensory processing, while parietal alpha rhythms are modulated by attentional effort and only indirectly affect sensory processing (Sokoliuk et al., 2019). Given the hypotheses we aimed to test in this paper, we discarded the parietal alpha rhythms from our analysis, even though our results show that SSD could be used to identify parietal alpha rhythms in resting-state in future studies. Third, brain size changes substantially during development and displays inter-individual variation, and we used a template brain averaged across ages 5–18. As such, our source localization procedure could have failed to properly localize components especially for the youngest and oldest children, whose brain sizes deviate most from the template brain used. Despite this concern, the distribution of identified components across age remained consistent with the participants’ age distribution (top right of Fig. 1A). We also chose to include premotor areas and use a slightly larger ROI for the sensorimotor region to compensate for possible imprecision in the source localization. Fourth, because our primary hypotheses focused on waveform shape differences across age and diagnoses, we focused on across-participants analyses and looked only at the component with highest SNR for each rhythm type (occipital alpha and sensorimotor mu). The large number of participants allowed us to make high-powered statistical inferences for our developmental research questions even with this limitation. That said, future studies should investigate whether these across-participant findings replicate within individuals in longitudinal data. These studies could use our spatial filtering approach to compare rhythms of the same type within participants, given that SSD is able to distinguish rhythms coming from the same cortical area in the same frequency band (Schaworonkow and Voytek, 2021a).

### Conclusion

Occipital alpha rhythms and sensorimotor mu rhythms are difficult to disentangle in sensor-space EEG due to volume conduction, with occipital alpha dominating even the central channels. Using data-driven spatial filters, we extracted time series for alpha and mu components with high alpha SNR from a large developmental dataset. Analysis of the time series of these alpha-band components replicated findings of lower frequency and higher amplitude for alpha than for mu. We demonstrated that mu waveforms are more asymmetrical than alpha waveforms. We extended previous findings of alpha-band rhythms undergoing large changes in frequency during development, showing that waveform shape measures also change substantially across development. While none of these resting-state changes appear to be associated with ADHD or autism in our dataset, characterizing differences across rhythm types remains vital for constraining generative models for alpha-band rhythms.

## Appendix A

**Figure A1:**
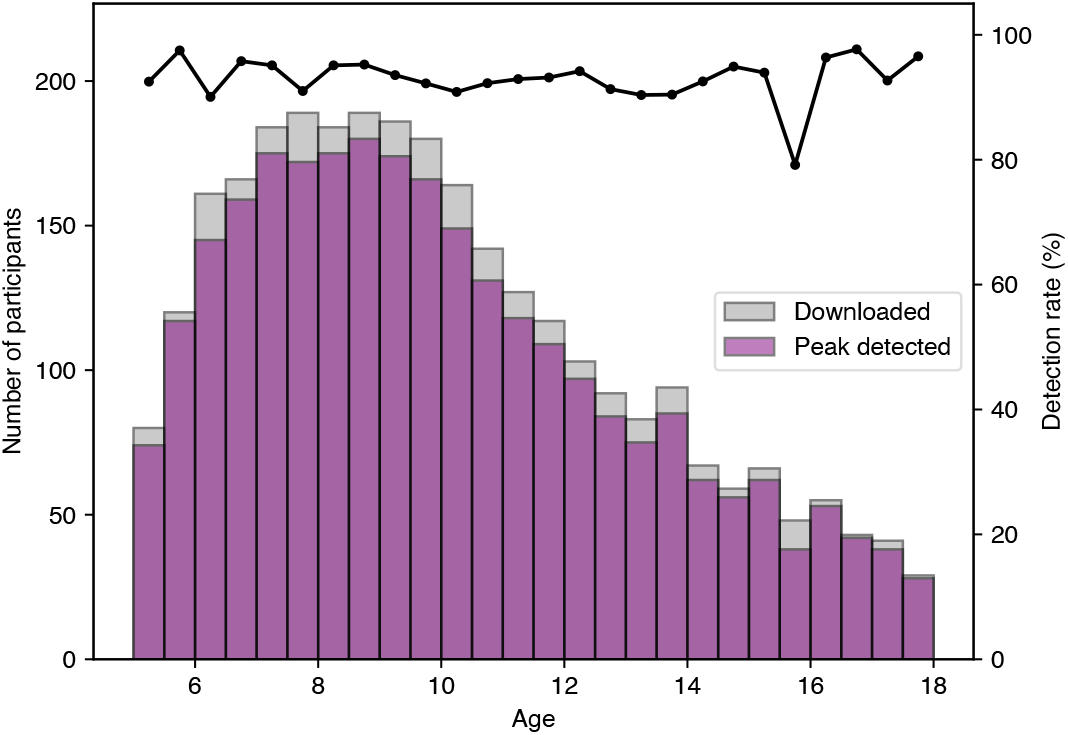
Alpha-band peak detection rate by age. Histogram of ages for participants with file downloaded and analyzed (gray) and with alpha peak detection by spectral parameterization (purple). The detection rate (black line) is the percentage of the participants for which an alpha peak was detected.

**Figure A2:**
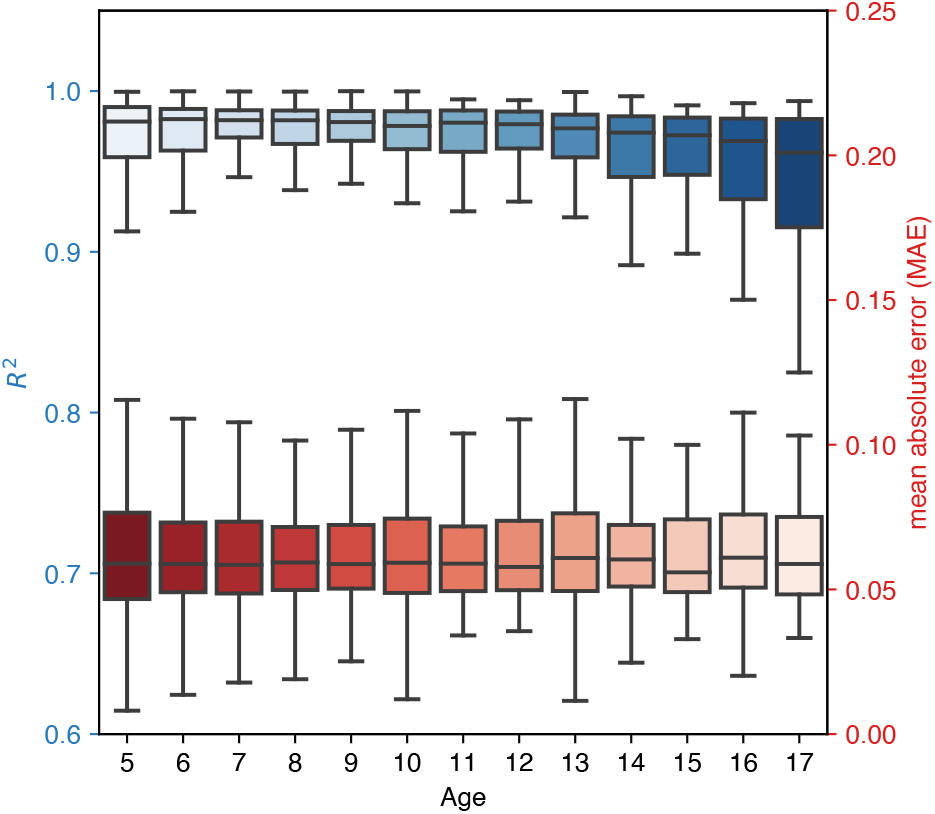
Goodness-of-fit metrics for spectral parameterization across development. *R*^2^ (blue) and mean absolute error (red) binned by year with lower bound of age shown, with the first bin being (5, 6].

**Figure A3:**
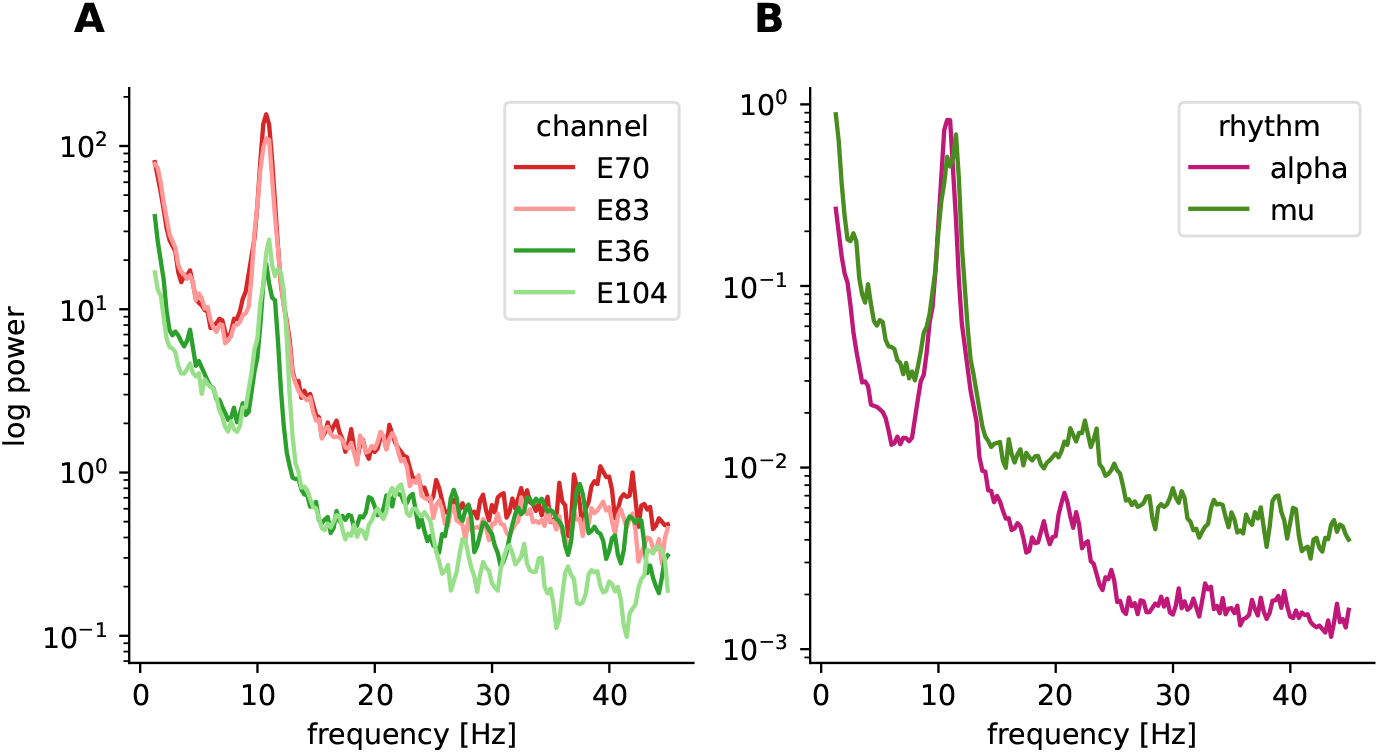
SSD removes high-frequency noise from power spectra. (A) Power spectra for two occipital channels (red) and two sensorimotor channels (green) from one participant. (B) Power spectra for alpha component (pink) and mu component (green) from the same participant.

## Appendix B

**Table B1:** Statistical testing for differences in waveform features between alpha and mu rhythms. t-values, Cohen’s d, and Bonferroni-corrected p-values across waveform features.

|  | frequency [Hz] | normalized amplitude | peak-trough asymmetry | rise-decay asymmetry |
| --- | --- | --- | --- | --- |
| within participants | $t=-2.965e+00$ | $t=1.436e+01$ | $t=-1.006e+01$ | $t=-4.476e+00$ |
| (paired t-test) | $p=2.514e-02$ | $p=4.492e-41$ | $p=1.406e-20$ | $p=7.509e-05$ |
| | $d=-1.685e-01$ | $d=6.972e-01$ | $d=-8.473e-01$ | $d=-2.947e-01$ |
| across participants | $t=-3.464e+00$ | $t=8.814e+00$ | $t=-9.003e+00$ | $t=-3.526e+00$ |
| (unpaired t-test) | $p=5.136e-03$ | $p=3.228e-15$ | $p=9.306e-16$ | $p=4.120e-03$ |
| | $d=-1.605e-01$ | $d=7.300e-01$ | $d=-8.602e-01$ | $d=-3.393e-01$ |

## Appendix C

**Figure C1:**
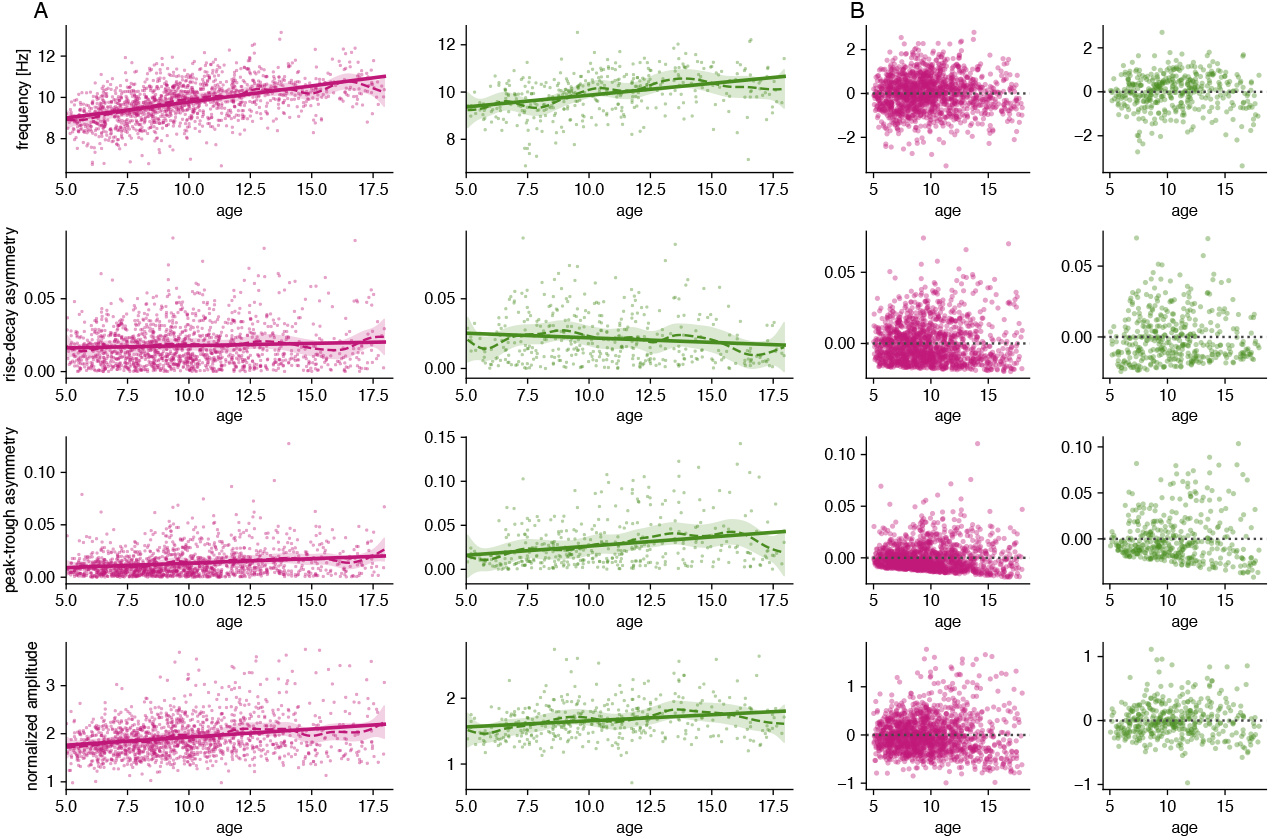
Assessing linear and nonlinear changes in waveform features with age. (A) Fit of waveform features with age for linear (solid line) and nonlinear (GAM) models (dashed line with 95% confidence interval) for alpha (pink, left) and mu (green, right) rhythms. (B) Residuals of linear model fits for waveform features with age for alpha (pink, left) and mu (green, right) rhythms.

**Figure C2:**
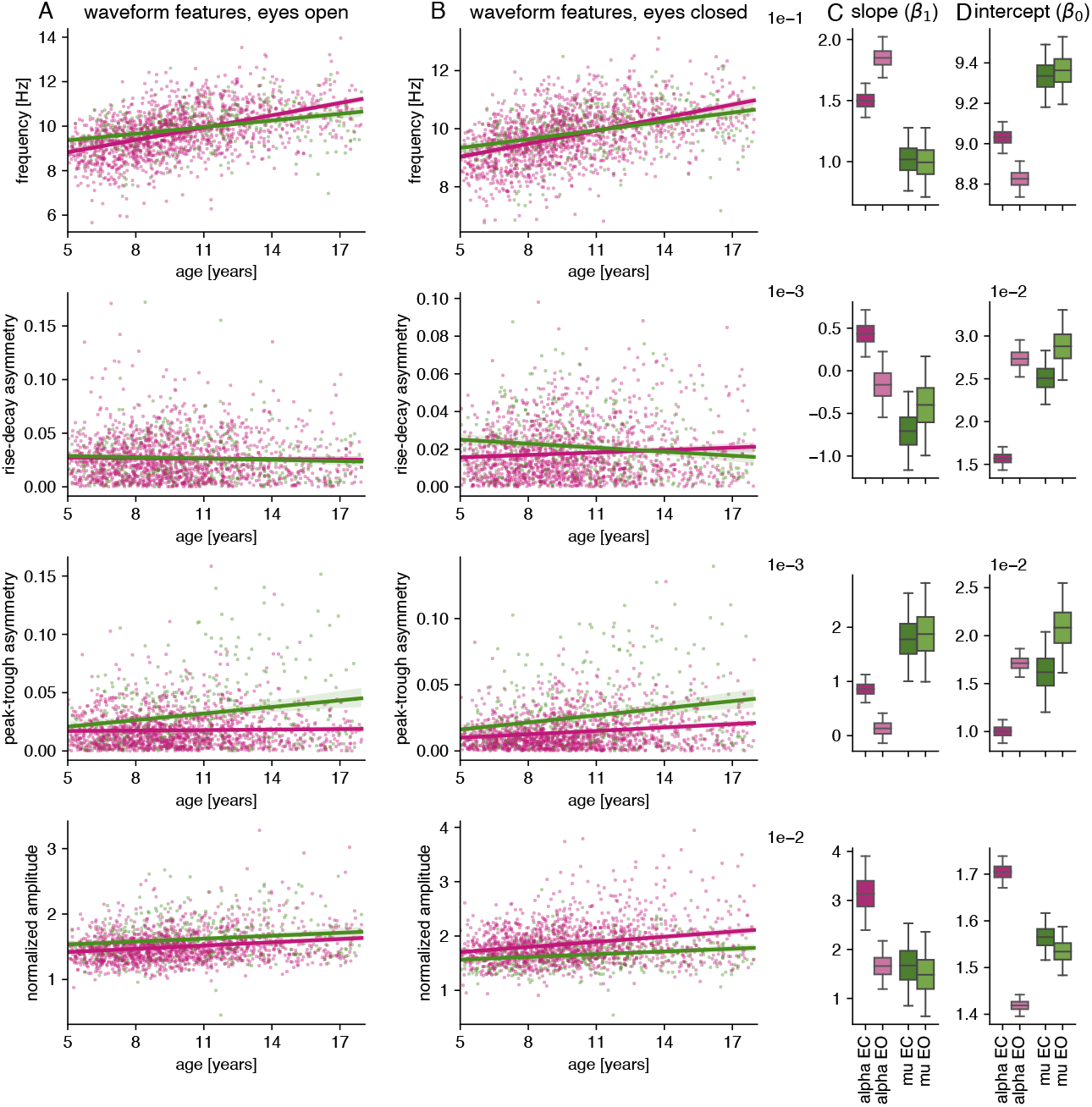
Alpha and mu rhythm waveform features across development for eyes open and eyes closed conditions. (A) Alpha (pink) and mu (green) waveform features (frequency, rise-decay and peak-trough asymmetry, normalized amplitude) by age across participants for the eyes closed condition. (B) same as (A) but for the eyes open condition. (C) Slope of linear model for alpha and mu rhythm for both eyes closed (EC) and eyes open (EO). Boxes denote interquartile range, and whiskers denote 95% confidence interval of bootstrapped feature values (10,000 iterations). (D) Intercept of linear model for alpha and mu rhythm for both EC and EO. Boxes denote interquartile range, and whiskers denote 95% confidence interval of bootstrapped feature values (10,000 iterations).

**Figure C3:**
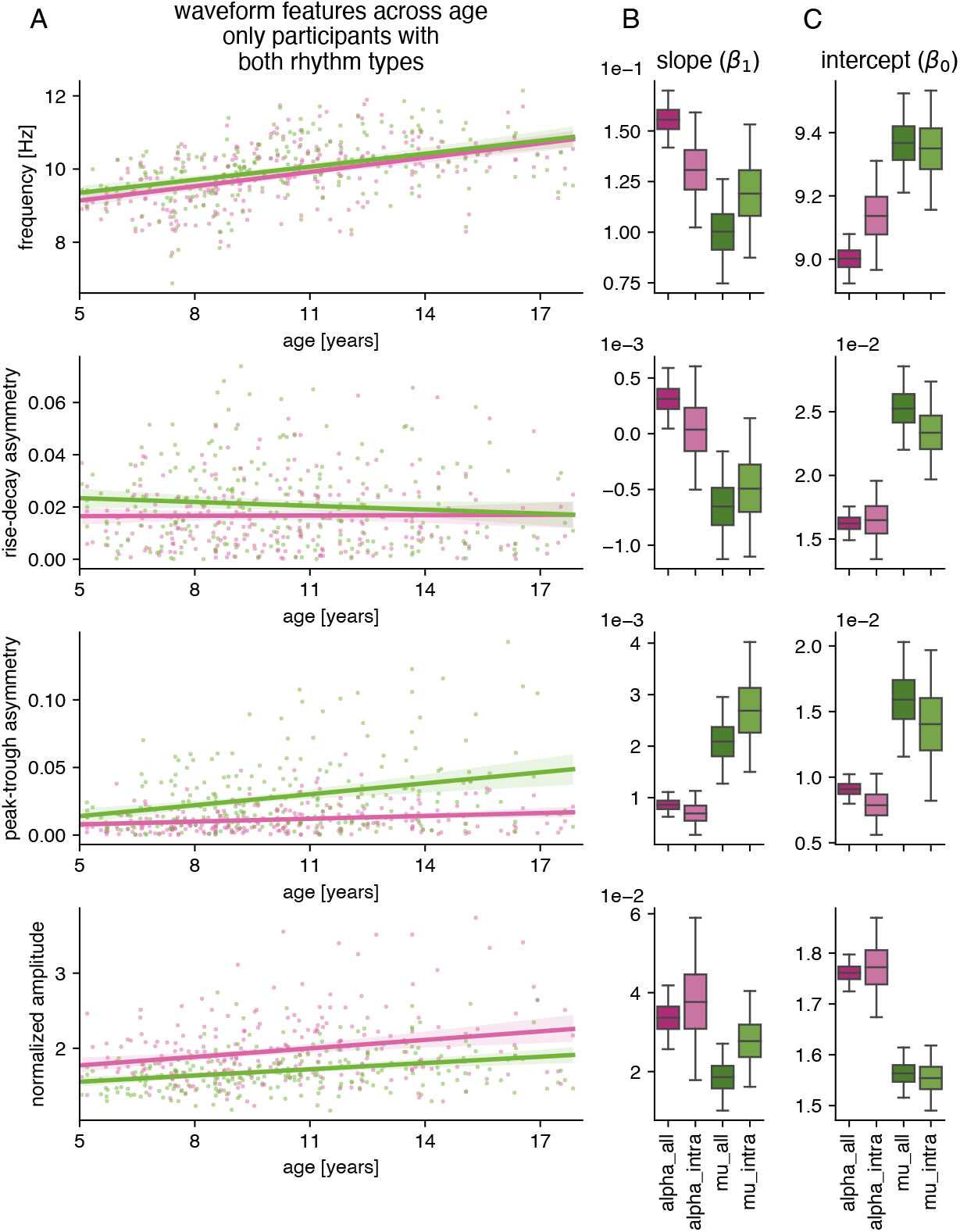
Waveform features across age, for participants with both rhythm types (N=239). (A) Alpha (pink) and mu (green) waveform features (frequency, rise-decay and peak-trough asymmetry, normalized amplitude) by age across participants who have both alpha and mu rhythms. (B) Slope of linear model for alpha and mu rhythm for all participants versus slopes for participants with both rhythm types (light pink and light green colors). Boxes denote interquartile range, and whiskers denote 95% confidence interval of bootstrapped feature values (10,000 iterations). (C) Intercept of linear model for alpha and mu rhythm for all participants versus intercepts for participants with both rhythm types. Boxes denote interquartile range, and whiskers denote 95% confidence interval of bootstrapped feature values, 10,000 iterations. Both slopes and intercepts for participants with both rhythm types are consistent with those across all participants.

## Appendix D

**Figure D1:**
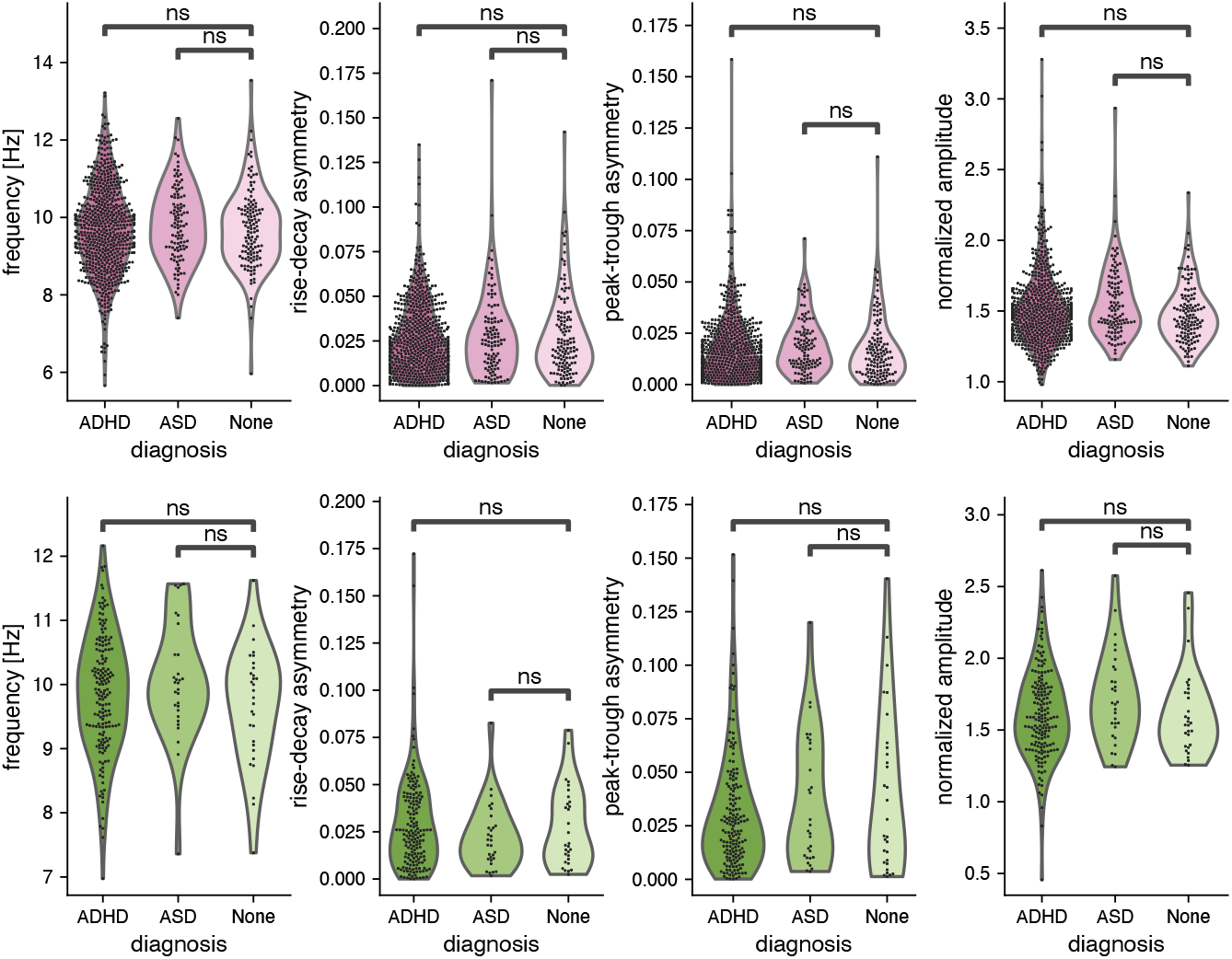
Waveform features with eyes open across ADHD and ASD diagnoses compared to typically developing participants. (A) Alpha rhythm components: waveform features (frequency, rise-decay and peak-trough asymmetry, normalized amplitude) with eyes open in participants who received an ADHD and ASD diagnosis, and for participants who did not receive a diagnosis (ADHD: N=639, ASD: N=129, no diagnosis N=109). (B) mu rhythm components: waveform features with eyes open in participants who received an ADHD and ASD diagnosis, and for participants without a diagnosis (ADHD: N=167, ASD: N=30, no diagnosis N=30). No statistically significant difference between the two disorder groups and typically developing participants was found for any waveform feature and rhythm type during eyes open.

**Figure D2:**
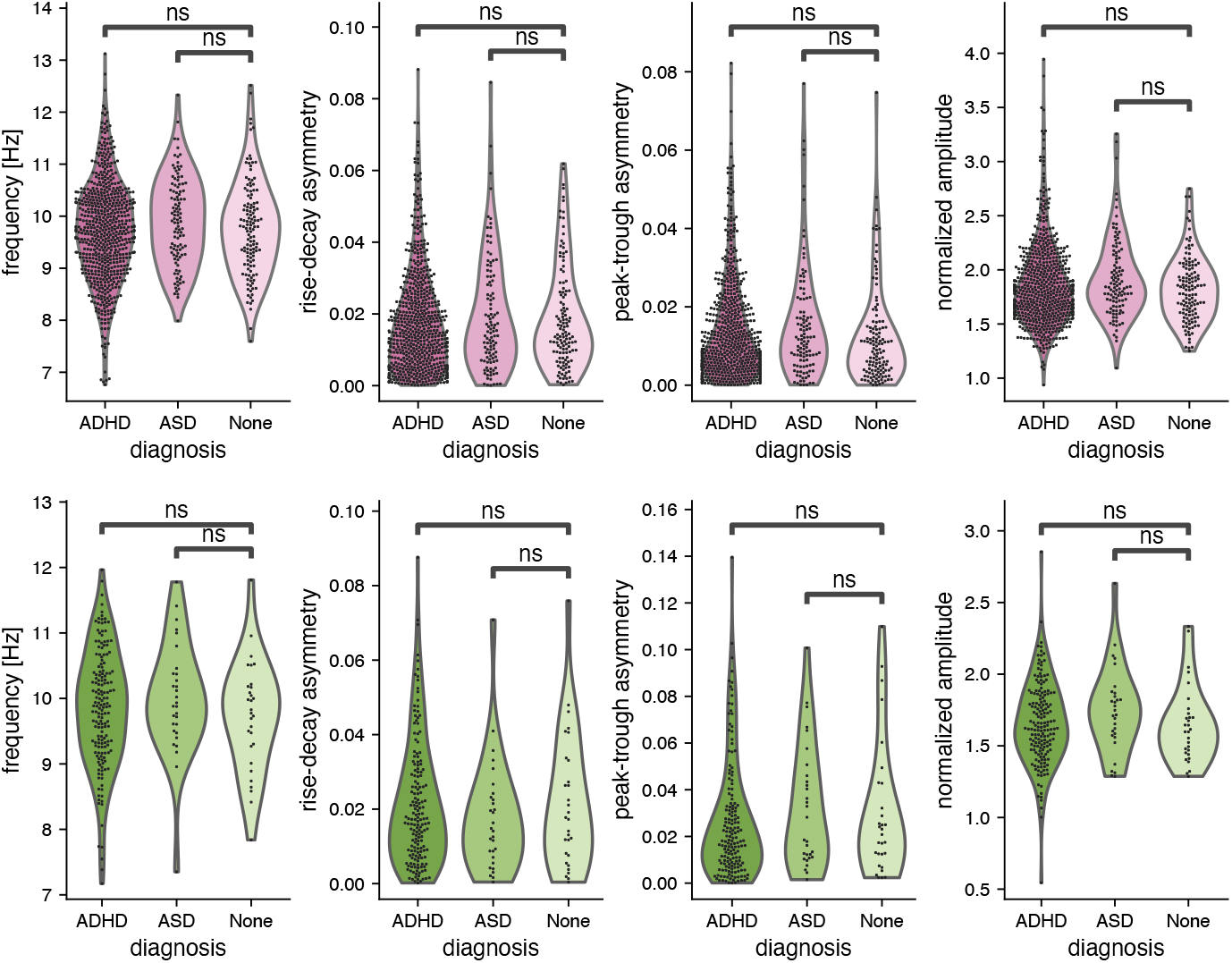
Waveform features with eyes closed across ADHD and ASD diagnoses compared to typically developing participants. (A) Alpha rhythm components: waveform features (frequency, rise-decay and peak-trough asymmetry, normalized amplitude) with eyes closed in participants who received an ADHD and ASD diagnosis, and for participants who did not receive a diagnosis (ADHD: N=639, ASD: N=129, no diagnosis N=109). (B) mu rhythm components: waveform features with eyes closed in participants who received an ADHD and ASD diagnosis, and for participants without a diagnosis (ADHD: N=167, ASD: N=30, no diagnosis N=30). No statistically significant difference between the two disorder groups and typically developing participants was found for any waveform feature and rhythm type during eyes closed.

**Figure D3:**
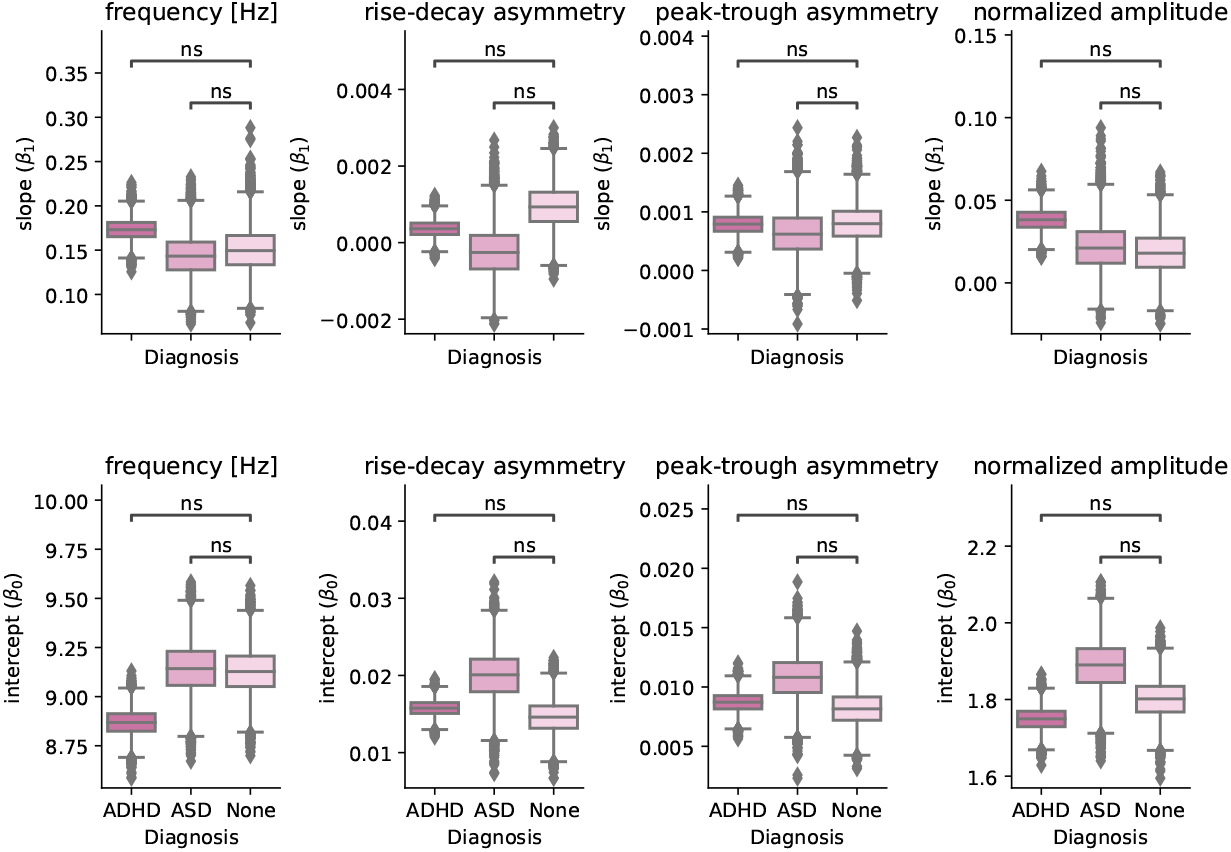
Alpha rhythm regression coefficient estimates using bootstrapping across diagnoses. Slope and intercept for three participant groups: participants who received an ADHD diagnosis, an ASD diagnosis, and participants who did not receive a diagnosis. Bootstrapping iterations: N=10,000.

**Figure D4:**
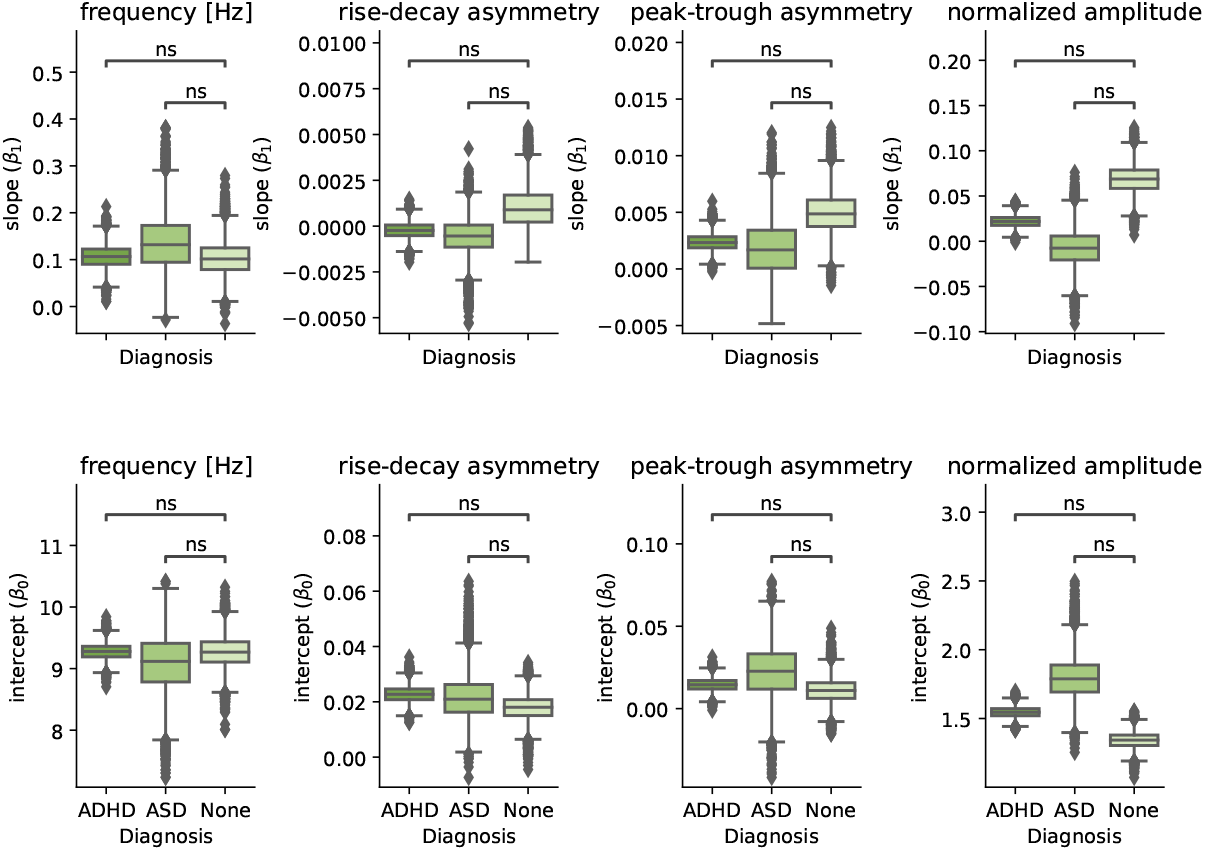
Mu rhythm regression coefficient estimates using bootstrapping across diagnoses. Same as Fig. D3, but for the mu rhythm.

**Table D1:** ANCOVA for each waveform feature across ADHD subtypes. The ANCOVA was performed across the following subtypes: Hyperactive/Impulsive, Inattentive, and Combined (both Hyperactive/Impulsive and Inattentive). There was no significant difference in waveform features between any of the subtypes and participants with no diagnosis, even before any correction for multiple comparisons was performed.

| rhythm | feature | Source | SS | DF | F | p-unc | np2 |
| --- | --- | --- | --- | --- | --- | --- | --- |
| Alpha | frequency [Hz] | Diagnosis | 3.079578 | 3 | 1.453295 | 0.226037 | 0.005689 |
| Alpha | rise-decay asymmetry | Diagnosis | 0.000453 | 3 | 0.765391 | 0.513629 | 0.003004 |
| Alpha | peak-trough asymmetry | Diagnosis | 0.000525 | 3 | 1.281284 | 0.279646 | 0.005019 |
| Alpha | normalized amplitude | Diagnosis | 0.134266 | 3 | 0.302939 | 0.823287 | 0.001191 |
| Mu | frequency [Hz] | Diagnosis | 3.178998 | 3 | 1.628719 | 0.184088 | 0.024817 |
| Mu | rise-decay asymmetry | Diagnosis | 0.000405 | 3 | 0.421879 | 0.737518 | 0.006549 |
| Mu | peak-trough asymmetry | Diagnosis | 0.002276 | 3 | 1.215632 | 0.305272 | 0.018640 |
| Mu | normalized amplitude | Diagnosis | 0.121427 | 3 | 0.525870 | 0.665016 | 0.008150 |

**Table D2:**
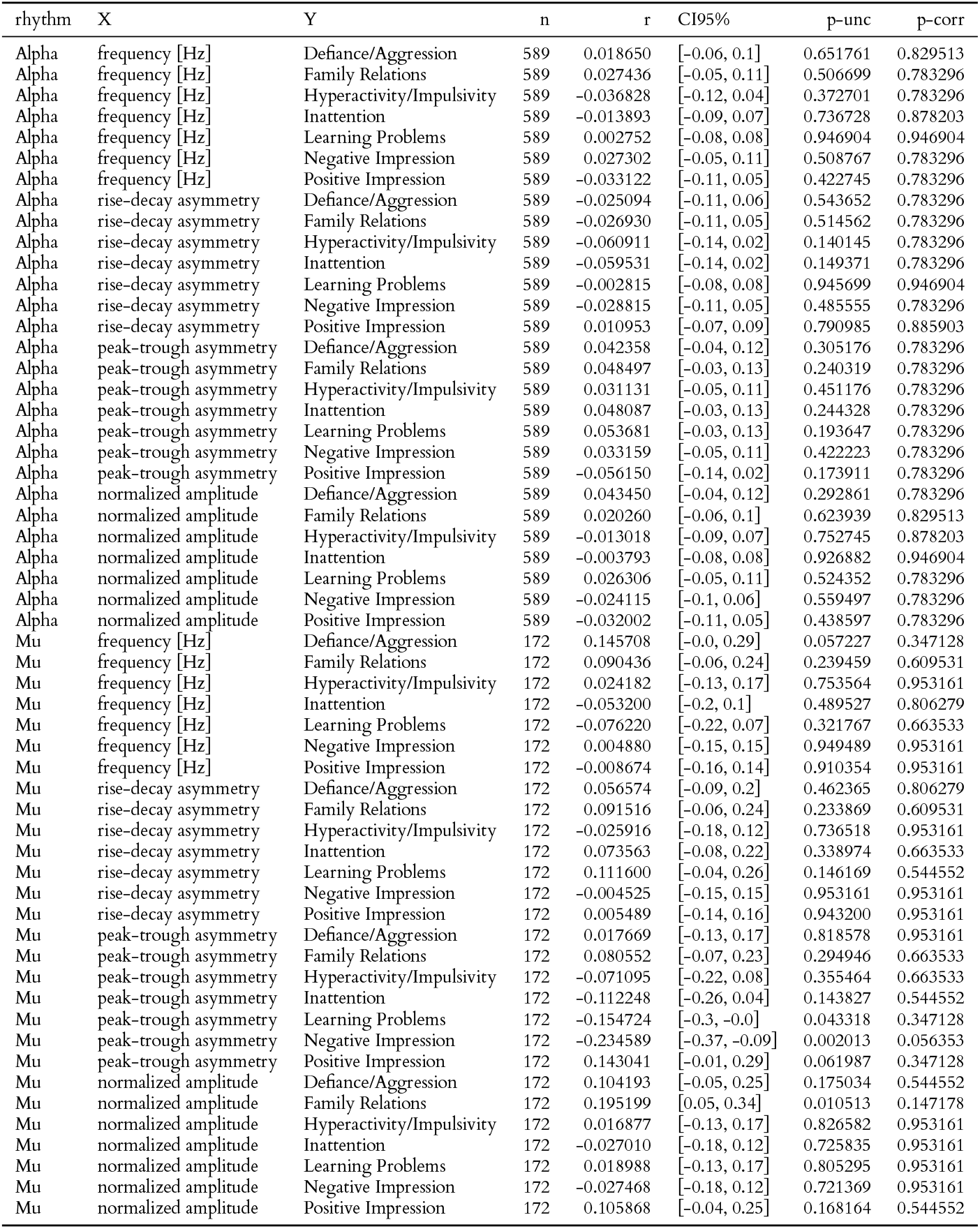
Correlation between Conners scores and waveform features. Correlation between different subscores of the Conners 3 - Self-Report and waveform features for all participants for which Conners scores are available.

**Table D3:** Correlation between Conners scores and waveform features, only ADHD group. Correlation between different subscores of the Conners 3 - Self-Report and waveform features for the ADHD group. See Table D3 for abbreviations.

| rhythm | X | Y | n | r | CI95% | p-unc | p-corr |
| --- | --- | --- | --- | --- | --- | --- | --- |
| Alpha | frequency [Hz] | Defiance/Aggression | 441 | 0.042072 | [-0.05, 0.14] | 0.378649 | 0.873179 |
| Alpha | frequency [Hz] | Family Relations | 441 | 0.076321 | [-0.02, 0.17] | 0.109887 | 0.675339 |
| Alpha | frequency [Hz] | Hyperactivity/Impulsivity | 441 | -0.009286 | [-0.1, 0.08] | 0.845988 | 0.906303 |
| Alpha | frequency [Hz] | Inattention | 441 | 0.033862 | [-0.06, 0.13] | 0.478652 | 0.873179 |
| Alpha | frequency [Hz] | Learning Problems | 441 | 0.015837 | [-0.08, 0.11] | 0.740439 | 0.906303 |
| Alpha | frequency [Hz] | Negative Impression | 441 | 0.028726 | [-0.06, 0.12] | 0.547859 | 0.873179 |
| Alpha | frequency [Hz] | Positive Impression | 441 | -0.033487 | [-0.13, 0.06] | 0.483530 | 0.873179 |
| Alpha | rise-decay asymmetry | Defiance/Aggression | 441 | -0.018054 | [-0.11, 0.08] | 0.705690 | 0.906303 |
| Alpha | rise-decay asymmetry | Family Relations | 441 | -0.031263 | [-0.12, 0.06] | 0.513069 | 0.873179 |
| Alpha | rise-decay asymmetry | Hyperactivity/Impulsivity | 441 | -0.047741 | [-0.14, 0.05] | 0.317726 | 0.873179 |
| Alpha | rise-decay asymmetry | Inattention | 441 | -0.019727 | [-0.11, 0.07] | 0.679854 | 0.906303 |
| Alpha | rise-decay asymmetry | Learning Problems | 441 | 0.031254 | [-0.06, 0.12] | 0.513188 | 0.873179 |
| Alpha | rise-decay asymmetry | Negative Impression | 441 | 0.005627 | [-0.09, 0.1] | 0.906303 | 0.906303 |
| Alpha | rise-decay asymmetry | Positive Impression | 441 | 0.007344 | [-0.09, 0.1] | 0.877916 | 0.906303 |
| Alpha | peak-trough asymmetry | Defiance/Aggression | 441 | 0.070797 | [-0.02, 0.16] | 0.138156 | 0.675339 |
| Alpha | peak-trough asymmetry | Family Relations | 441 | 0.068681 | [-0.02, 0.16] | 0.150359 | 0.675339 |
| Alpha | peak-trough asymmetry | Hyperactivity/Impulsivity | 441 | 0.068798 | [-0.02, 0.16] | 0.149666 | 0.675339 |
| Alpha | peak-trough asymmetry | Inattention | 441 | 0.040079 | [-0.05, 0.13] | 0.401669 | 0.873179 |
| Alpha | peak-trough asymmetry | Learning Problems | 441 | 0.077361 | [-0.02, 0.17] | 0.105116 | 0.675339 |
| Alpha | peak-trough asymmetry | Negative Impression | 441 | 0.008447 | [-0.09, 0.1] | 0.859752 | 0.906303 |
| Alpha | peak-trough asymmetry | Positive Impression | 441 | -0.066632 | [-0.16, 0.03] | 0.162940 | 0.675339 |
| Alpha | normalized amplitude | Defiance/Aggression | 441 | 0.062182 | [-0.03, 0.15] | 0.192954 | 0.675339 |
| Alpha | normalized amplitude | Family Relations | 441 | 0.027765 | [-0.07, 0.12] | 0.561329 | 0.873179 |
| Alpha | normalized amplitude | Hyperactivity/Impulsivity | 441 | -0.009206 | [-0.1, 0.08] | 0.847299 | 0.906303 |
| Alpha | normalized amplitude | Inattention | 441 | -0.023765 | [-0.12, 0.07] | 0.619079 | 0.906303 |
| Alpha | normalized amplitude | Learning Problems | 441 | 0.035752 | [-0.06, 0.13] | 0.454428 | 0.873179 |
| Alpha | normalized amplitude | Negative Impression | 441 | -0.063099 | [-0.16, 0.03] | 0.186460 | 0.675339 |
| Alpha | normalized amplitude | Positive Impression | 441 | -0.012462 | [-0.11, 0.08] | 0.794352 | 0.906303 |
| Mu | frequency [Hz] | Defiance/Aggression | 128 | 0.179917 | [0.01, 0.34] | 0.042963 | 0.449300 |
| Mu | frequency [Hz] | Family Relations | 128 | 0.161064 | [-0.01, 0.33] | 0.070453 | 0.449300 |
| Mu | frequency [Hz] | Hyperactivity/Impulsivity | 128 | 0.000673 | [-0.17, 0.17] | 0.994007 | 0.994007 |
| Mu | frequency [Hz] | Inattention | 128 | -0.062326 | [-0.23, 0.11] | 0.486362 | 0.716744 |
| Mu | frequency [Hz] | Learning Problems | 128 | -0.055451 | [-0.23, 0.12] | 0.535782 | 0.750095 |
| Mu | frequency [Hz] | Negative Impression | 128 | 0.006848 | [-0.17, 0.18] | 0.939091 | 0.994007 |
| Mu | frequency [Hz] | Positive Impression | 128 | -0.004934 | [-0.18, 0.17] | 0.956091 | 0.994007 |
| Mu | rise-decay asymmetry | Defiance/Aggression | 128 | 0.073236 | [-0.1, 0.24] | 0.413207 | 0.642766 |
| Mu | rise-decay asymmetry | Family Relations | 128 | 0.145279 | [-0.03, 0.31] | 0.103169 | 0.449300 |
| Mu | rise-decay asymmetry | Hyperactivity/Impulsivity | 128 | -0.047378 | [-0.22, 0.13] | 0.596845 | 0.759621 |
| Mu | rise-decay asymmetry | Inattention | 128 | 0.048717 | [-0.13, 0.22] | 0.586506 | 0.759621 |
| Mu | rise-decay asymmetry | Learning Problems | 128 | 0.122858 | [-0.05, 0.29] | 0.168799 | 0.449300 |
| Mu | rise-decay asymmetry | Negative Impression | 128 | 0.078933 | [-0.1, 0.25] | 0.377716 | 0.642766 |
| Mu | rise-decay asymmetry | Positive Impression | 128 | 0.124824 | [-0.05, 0.29] | 0.162035 | 0.449300 |
| Mu | peak-trough asymmetry | Defiance/Aggression | 128 | 0.032824 | [-0.14, 0.21] | 0.714107 | 0.833125 |
| Mu | peak-trough asymmetry | Family Relations | 128 | 0.120688 | [-0.05, 0.29] | 0.176511 | 0.449300 |
| Mu | peak-trough asymmetry | Hyperactivity/Impulsivity | 128 | -0.108042 | [-0.28, 0.07] | 0.226634 | 0.528812 |
| Mu | peak-trough asymmetry | Inattention | 128 | -0.088472 | [-0.26, 0.09] | 0.322607 | 0.642766 |
| Mu | peak-trough asymmetry | Learning Problems | 128 | -0.087755 | [-0.26, 0.09] | 0.326560 | 0.642766 |
| Mu | peak-trough asymmetry | Negative Impression | 128 | -0.139189 | [-0.31, 0.04] | 0.118599 | 0.449300 |
| Mu | peak-trough asymmetry | Positive Impression | 128 | 0.174109 | [-0.0, 0.34] | 0.050267 | 0.449300 |
| Mu | normalized amplitude | Defiance/Aggression | 128 | 0.121244 | [-0.05, 0.29] | 0.174509 | 0.449300 |
| Mu | normalized amplitude | Family Relations | 128 | 0.290474 | [0.12, 0.44] | 0.000924 | 0.025859 |
| Mu | normalized amplitude | Hyperactivity/Impulsivity | 128 | -0.001875 | [-0.18, 0.17] | 0.983306 | 0.994007 |
| Mu | normalized amplitude | Inattention | 128 | 0.039592 | [-0.14, 0.21] | 0.658528 | 0.801687 |
| Mu | normalized amplitude | Learning Problems | 128 | 0.142066 | [-0.03, 0.31] | 0.111102 | 0.449300 |
| Mu | normalized amplitude | Negative Impression | 128 | 0.075043 | [-0.1, 0.25] | 0.401740 | 0.642766 |
| Mu | normalized amplitude | Positive Impression | 128 | 0.074188 | [-0.1, 0.25] | 0.407142 | 0.642766 |

**Table D4:**
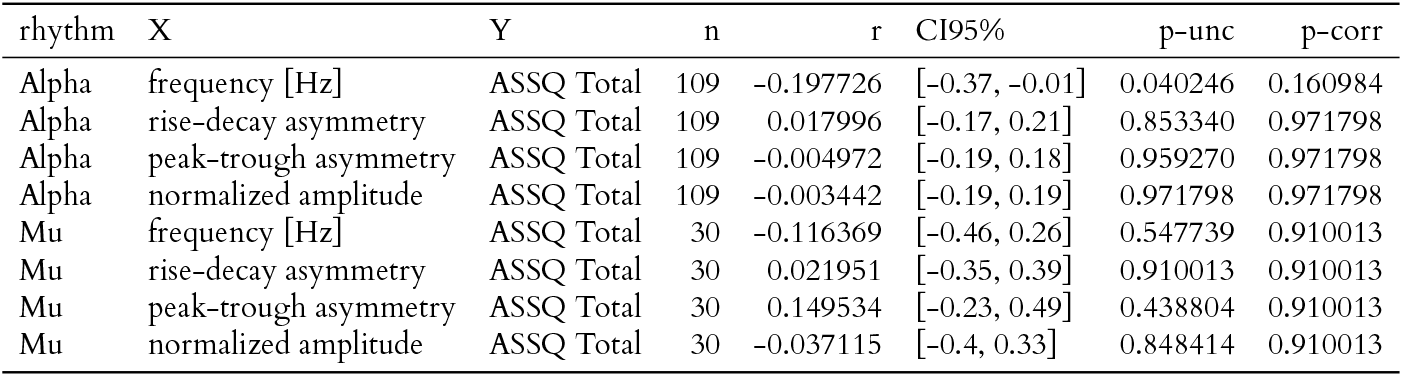
Correlation between ASSQ scores and waveform features, only ASD group. No significant correlation was found between any of the waveform features and the ASSQ scores.

| rhythm | X | Y | n | r | CI95% | p-unc | p-corr |
| --- | --- | --- | --- | --- | --- | --- | --- |
| Alpha | frequency [Hz] | ASSQ Total | 109 | -0.197726 | [-0.37, -0.01] | 0.040246 | 0.160984 |
| Alpha | rise-decay asymmetry | ASSQ Total | 109 | 0.017996 | [-0.17, 0.21] | 0.853340 | 0.971798 |
| Alpha | peak-trough asymmetry | ASSQ Total | 109 | -0.004972 | [-0.19, 0.18] | 0.959270 | 0.971798 |
| Alpha | normalized amplitude | ASSQ Total | 109 | -0.003442 | [-0.19, 0.19] | 0.971798 | 0.971798 |
| Mu | frequency [Hz] | ASSQ Total | 30 | -0.116369 | [-0.46, 0.26] | 0.547739 | 0.910013 |
| Mu | rise-decay asymmetry | ASSQ Total | 30 | 0.021951 | [-0.35, 0.39] | 0.910013 | 0.910013 |
| Mu | peak-trough asymmetry | ASSQ Total | 30 | 0.149534 | [-0.23, 0.49] | 0.438804 | 0.910013 |
| Mu | normalized amplitude | ASSQ Total | 30 | -0.037115 | [-0.4, 0.33] | 0.848414 | 0.910013 |

**Table D5:** Correlation between ASSQ scores and waveform features, all participants for which ASSQ is available. No significant correlation was found between any of the waveform features and the ASSQ scores.

| rhythm | X | Y | n | r | CI95% | p-unc | p-corr |
| --- | --- | --- | --- | --- | --- | --- | --- |
| Alpha | frequency [Hz] | ASSQ Total | 873 | 0.025724 | [-0.04, 0.09] | 0.448055 | 0.794258 |
| Alpha | rise-decay asymmetry | ASSQ Total | 873 | -0.008844 | [-0.08, 0.06] | 0.794258 | 0.794258 |
| Alpha | peak-trough asymmetry | ASSQ Total | 873 | 0.017391 | [-0.05, 0.08] | 0.608063 | 0.794258 |
| Alpha | normalized amplitude | ASSQ Total | 873 | 0.015952 | [-0.05, 0.08] | 0.638065 | 0.794258 |
| Mu | frequency [Hz] | ASSQ Total | 225 | 0.024818 | [-0.11, 0.16] | 0.711813 | 0.889591 |
| Mu | rise-decay asymmetry | ASSQ Total | 225 | -0.074651 | [-0.2, 0.06] | 0.265892 | 0.889591 |
| Mu | peak-trough asymmetry | ASSQ Total | 225 | 0.050103 | [-0.08, 0.18] | 0.455580 | 0.889591 |
| Mu | normalized amplitude | ASSQ Total | 225 | 0.009327 | [-0.12, 0.14] | 0.889591 | 0.889591 |

**Table D6:** ANCOVA for each waveform feature across secondary diagnoses. ANCOVA statistics to test whether there is difference in waveform features after separating participants with both ADHD and ASD diagnoses from participants with only ADHD or ASD.

| Rhythm | Feature | F | np2 | p-unc | p-corr |
| --- | --- | --- | --- | --- | --- |
| Alpha | frequency [Hz] | 1.836953 | 0.006287 | 0.138839 | 0.828708 |
| Alpha | rise-decay asymmetry | 1.022492 | 0.003509 | 0.381886 | 0.828708 |
| Alpha | peak-trough asymmetry | 0.331548 | 0.001141 | 0.802548 | 0.937907 |
| Alpha | normalized amplitude | 0.953035 | 0.003272 | 0.414354 | 0.828708 |
| Mu | frequency [Hz] | 0.136906 | 0.001847 | 0.937907 | 0.937907 |
| Mu | rise-decay asymmetry | 0.247669 | 0.003336 | 0.862943 | 0.937907 |
| Mu | peak-trough asymmetry | 1.452749 | 0.019254 | 0.228374 | 0.828708 |
| Mu | normalized amplitude | 0.446848 | 0.006002 | 0.719772 | 0.937907 |

**Table D7:** Relationship between WISC scores and diagnosis. Results from ANCOVA for WISC scores with main factor of diagnosis with age as a covariate.

| WISC score | Source | F | p-unc | np2 |
| --- | --- | --- | --- | --- |
| Verbal Comprehension Index | Diagnosis | 16.408737 | 9.853858e-08 | 0.033256 |
| Visual Spatial Index | Diagnosis | 7.295580 | 7.170606e-04 | 0.015064 |
| Processing Speed Index | Diagnosis | 19.862316 | 3.540166e-09 | 0.040056 |
| Working Memory Index | Diagnosis | 20.406914 | 2.097635e-09 | 0.041027 |
| Fluid Reasoning Index | Diagnosis | 9.386124 | 9.189387e-05 | 0.019318 |
| FSIQ | Diagnosis | 20.991018 | 1.198413e-09 | 0.042151 |

**Table D8:** Linear model with age and WISC scores as covariates. We fit the following linear model for each waveform feature: *WF* ~ *Age* + *Diagnosis* + *WISC FSIQ scores*. All waveform features remain statistically nonsignificant.

| feature | rhythm | Coef. | Std.Err. | P> \|t\| | [0.025 | 0.975] |
| --- | --- | --- | --- | --- | --- | --- |
| frequency [Hz] | Alpha | 0.002084 | 0.001951 | 0.285789 | -0.001746 | 0.005914 |
| rise-decay asymmetry | Alpha | -0.000005 | 0.000032 | 0.884289 | -0.000068 | 0.000059 |
| peak-trough asymmetry | Alpha | 0.000021 | 0.000028 | 0.456528 | -0.000034 | 0.000075 |
| normalized amplitude | Alpha | 0.001160 | 0.000887 | 0.191174 | -0.000581 | 0.002901 |
| frequency [Hz] | Mu | 0.003592 | 0.003918 | 0.360438 | -0.004136 | 0.011320 |
| rise-decay asymmetry | Mu | -0.000024 | 0.000085 | 0.778237 | -0.000191 | 0.000144 |
| peak-trough asymmetry | Mu | 0.000016 | 0.000118 | 0.895043 | -0.000218 | 0.000249 |
| normalized amplitude | Mu | 0.001272 | 0.001342 | 0.344192 | -0.001374 | 0.003919 |

**Figure D5:**
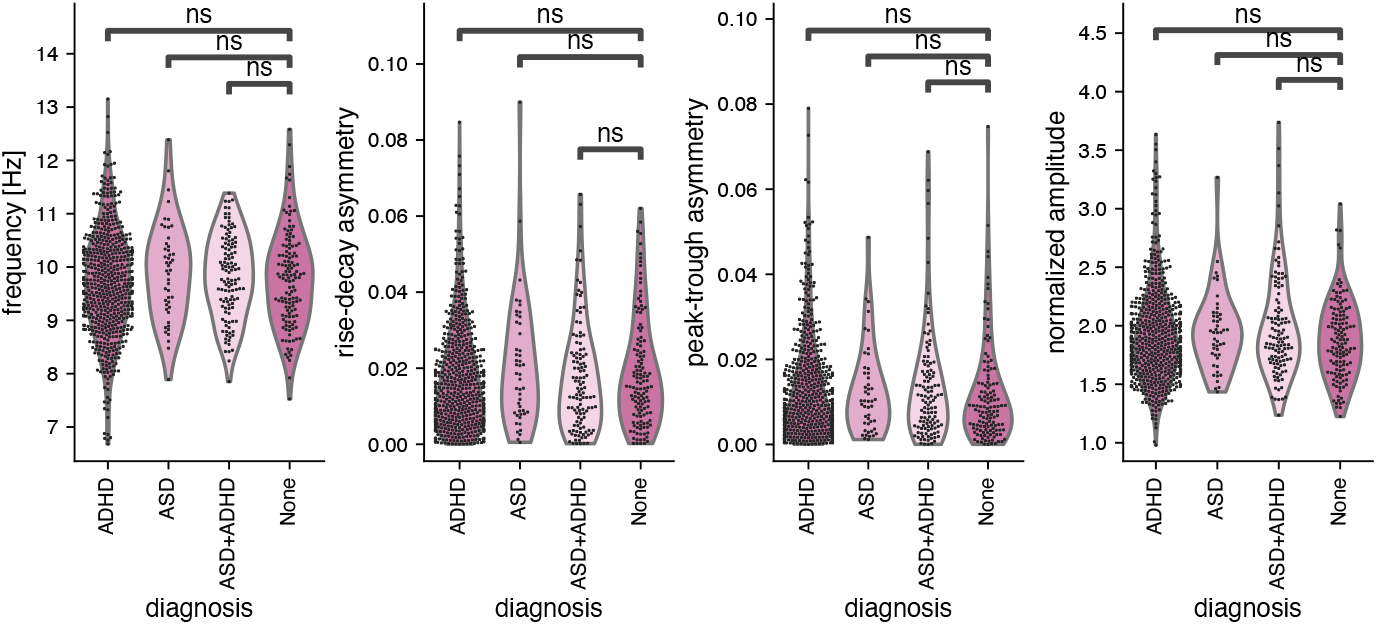
Alpha waveform features across diagnoses, ADHD+ASD group separately. There is no significant difference in alpha waveform features between participants with both ADHD and ASD, ADHD only, ASD only, and participants with no diagnosis.

**Figure D6:**
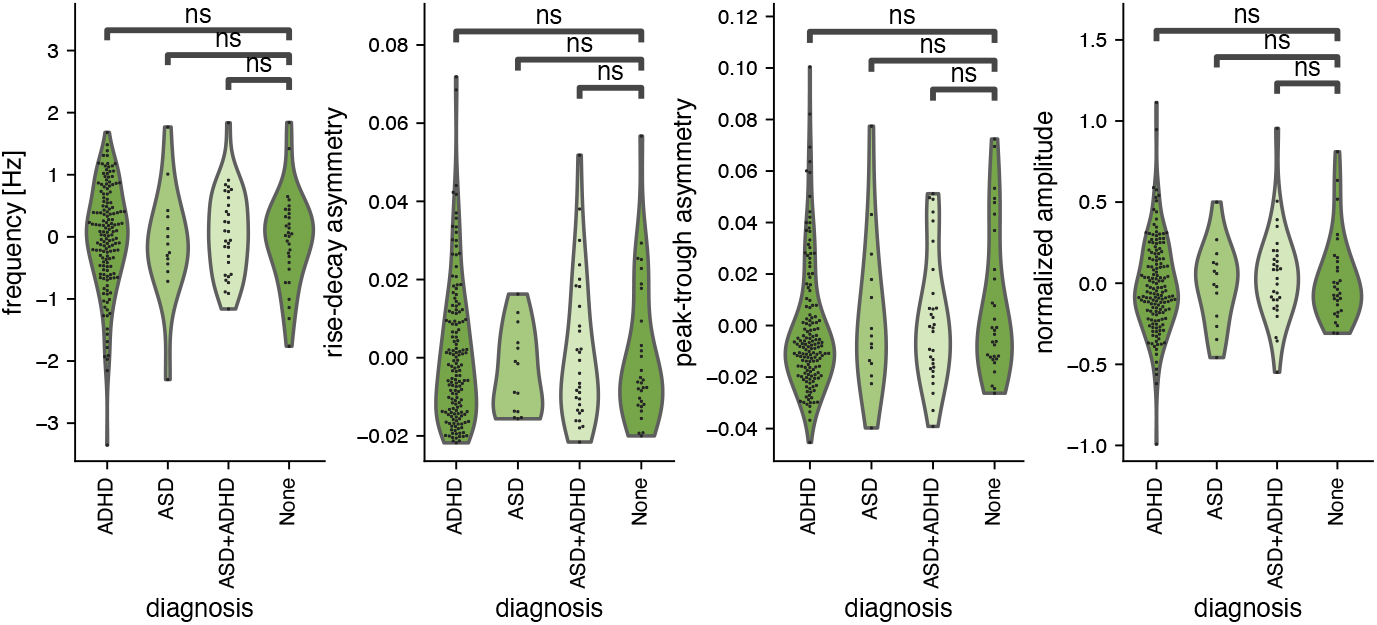
Mu waveform features across diagnoses, ADHD+ASD group separately. There is no significant difference in mu waveform features between participants with both ADHD and ASD, ADHD only, ASD only, and participants with no diagnosis.

**Figure D7:**
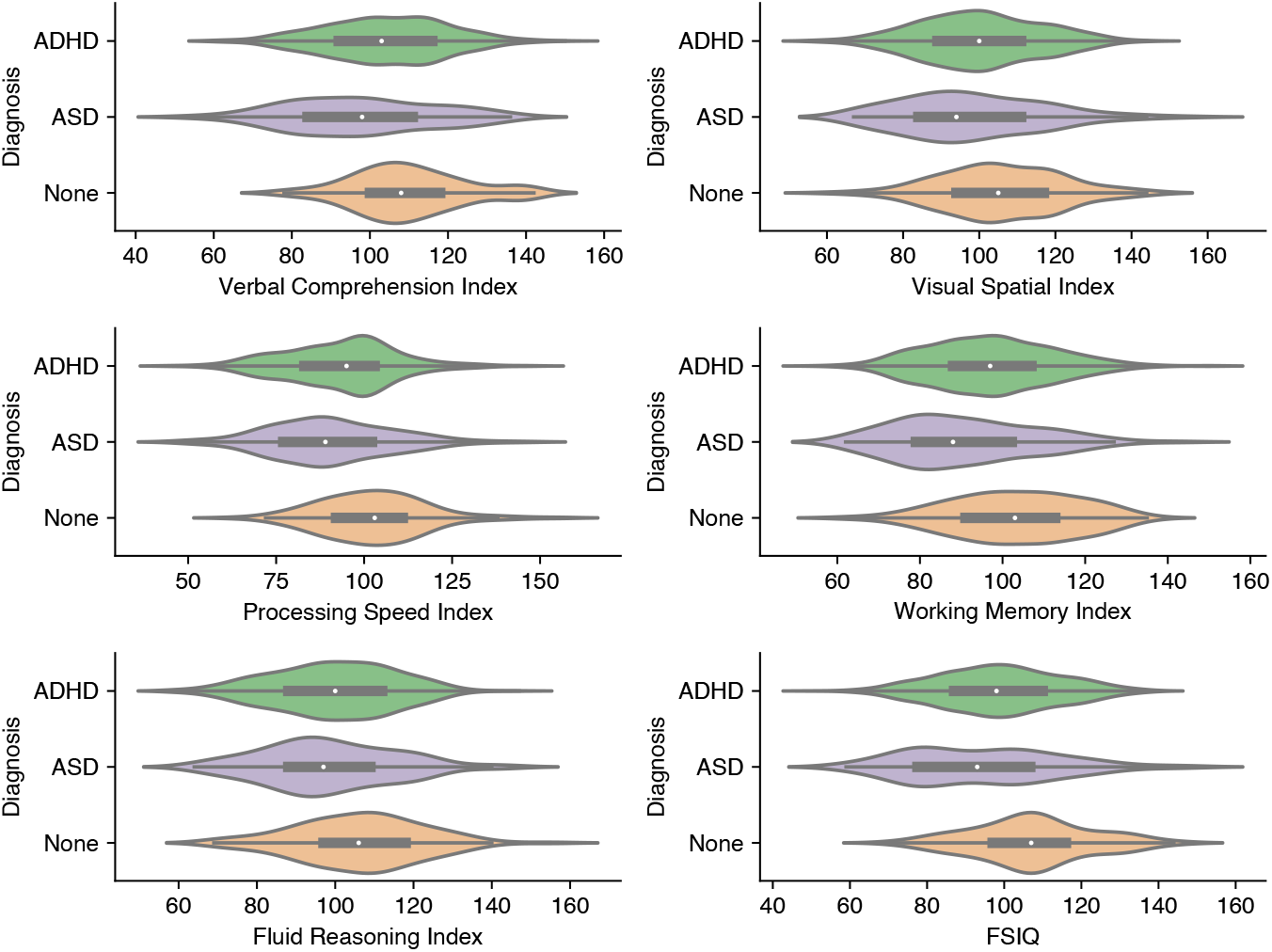
WISC-V scores across diagnoses. Scores for Wechsler Intelligence Scale for Children (Fifth Edition) across diagnoses. All six scores show significant effect of diagnosis.

## Data and code availability

The data are available through the scientific data portal of the Healthy Brain Network dataset from the Child Mind Institute at the following link: https://fcon_1000.projects.nitrc.org/indi/cmi_healthy_brain_network/. EEG data is distributed under the Creative Commons, Attribution Non-Commercial Share Alike License. Access to the phenotypic data, including the diagnostic data used in this manuscript, requires completion of the Data Usage Agreement (DUA) available at the following link: https://fcon_1000.projects.nitrc.org/indi/cmi_healthy_brain_network/Pheno_Access.html. All the necessary code to reproduce the analysis and figures of this manuscript are available at the following GitHub repository: https://github.com/nschawor/eeg-mu-alpha-development.

## Author contributions

A.B.: Conceptualization, Methodology, Software, Formal Analysis, Visualization, Writing-Original Draft Preparation. B.V.: Conceptualization, Writing - Review & Editing, Supervision, Resources, Project Administration, Funding acquisition. N.S.: Conceptualization, Methodology, Software, Formal Analysis, Visualization, Writing-Original Draft Preparation, Supervision.

## Funding

This work was supported by funding from the European Union’s Horizon Europe research and innovation programme to N.S. under the Marie Skłodowska-Curie Postdoctoral Fellowship grant agreement No. 101062497 and from the NIH National Institute of General Medical Sciences grant R01GM134363-01 to B.V.

## Declaration of competing interests

The authors declare no competing interests.

## Acknowledgements

This manuscript was prepared using a limited access dataset obtained from the Child Mind Institute Biobank, Healthy Brain Network. This manuscript reflects the views of the authors and does not necessarily reflect the opinions or views of the Child Mind Institute.

## Notes

http://fcon_1000.projects.nitrc.org/indi/cmi_healthy_brain_network/sharing_neuro.html

### Competing Interest Statement

The authors have declared no competing interest.

### Summary of Updates

We changed the title following peer review

## References

Aird, RB and Gastaut, Y (1959). “Occipital and posterior electroencephalographic ryhthms”. In: Electroencephalography and clinical neurophysiology 11.4, pp. 637–656.

Alexander, LM et al. (2017). “An open resource for transdiagnostic research in pediatric mental health and learning disorders”. In: Scientific Data 4.1, pp. 170–181. DOI: 10.1038/sdata.2017.181.

Baio, J (2018). “Prevalence of autism spectrum disorder among children aged 8 years—autism and developmental disabilities monitoring network, 11 sites, United States, 2014”. In: MMWR. Surveillance Summaries 67.

Benjamini, Y and Hochberg, Y (1995). “Controlling the false discovery rate: a practical and powerful approach to multiple testing”. In: Journal of the Royal statistical society: series B (Methodological) 57.1, pp. 289–300.

Berger, H (1929). “Über das Elektrenkephalogramm des Menschen”. In: Archiv für Psychiatrie und Nervenkrankheiten.

Bethlehem, RAI et al. (2022). “Brain charts for the human lifespan”. In: Nature 604.7906, pp. 525–533. DOI: 10.1038/s41586-022-04554-y.

Bitzenhofer, SH, Pöpplau, JA, and Hanganu-Opatz, I (2020). “Gamma activity accelerates during prefrontal development”. en. In: eLife 9, e56795. DOI: 10.7554/eLife.56795.

Caffarra, S et al. (2024). “Development of the Alpha Rhythm Is Linked to Visual White Matter Pathways and Visual Detection Performance”. In: Journal of Neuroscience 44.6. DOI: 10.1523/JNEUROSCI.0684-23.2023.

Caveness, W (1962). Atlas of Electroencephalography in the Developing Monkey Macaca Mulatta. Addison-Wesley.

Cellier, D, Riddle, J, Petersen, I, and Hwang, K (2021). “The development of theta and alpha neural oscillations from ages 3 to 24 years”. In: Developmental cognitive neuroscience 50, p. 100969.

Cole, SR, van der Meij, R, Peterson, EJ, Hemptinne, C de, Starr, PA, and Voytek, B (2017). “Nonsinusoidal Beta Oscillations Reflect Cortical Pathophysiology in Parkinson’s Disease”. In: The Journal of Neuroscience 37.18, pp. 4830–4840. DOI: 10.1523/JNEUROSCI.2208-16.2017.

Cole, SR and Voytek, B (2017). “Brain Oscillations and the Importance of Waveform Shape”. In: Trends in Cognitive Sciences 21.2, pp. 137–149. DOI: 10.1016/j.tics.2016.12.008.

Cole, SR and Voytek, B (2019). “Cycle-by-cycle analysis of neural oscillations”. In: Journal of Neurophysiology 122.2, pp. 849–861. DOI: 10.1152/jn.00273.2019.

Danielson, ML, Bitsko, RH, Ghandour, RM, Holbrook, JR, Kogan, MD, and Blumberg, SJ (2018). “Prevalence of parent-reported ADHD diagnosis and associated treatment among US children and adolescents, 2016”. In: Journal of Clinical Child & Adolescent Psychology 47.2, pp. 199–212.

Dede, AJ, Xiao, W, Vaci, N, Cohen, MX, and Milne, E (2023). “Lack of univariate, clinically-relevant biomarkers of autism in resting state EEG: a study of 776 participants”. In: medRxiv. DOI: 10.1101/2023.05.21.23290300.

Donoghue, T, Haller, M, et al. (2020). “Parameterizing neural power spectra into periodic and aperiodic components”. In: Nature Neuroscience 23.12, pp. 1655–1665. DOI: 10.1038/s41593-020-00744-x.

Donoghue, T, Schaworonkow, N, and Voytek, B (2021). “Methodological considerations for studying neural oscillations”. In: European Journal of Neuroscience. ISSN: 0953-816X, 1460-9568. DOI: 10.1111/ejn.15361.

Dumas, G, Soussignan, R, Hugueville, L, Martinerie, J, and Nadel, J (2014). “Revisiting mu suppression in autism spectrum disorder”. In: Brain research 1585, pp. 108–119.

Ede, F van (2018). “Mnemonic and attentional roles for states of attenuated alpha oscillations in perceptual working memory: a review”. In: The European Journal of Neuroscience 48.7, pp. 2509–2515. DOI: 10.1111/ejn.13759.

Fischl, B (2012). “FreeSurfer”. In: NeuroImage 62.2, pp. 774–781. DOI: 10.1016/j.neuroimage.2012.01.021.

Fonov, V, Evans, AC, Botteron, K, Almli, CR, McKinstry, RC, and Collins, DL (2011). “Unbiased average age-appropriate atlases for pediatric studies”. In: NeuroImage 54.1, pp. 313–327. DOI: 10.1016/j.neuroimage.2010.07.033.

Foster, JJ, Sutterer, DW, Serences, JT, Vogel, EK, and Awh, E (2016). “The topography of alpha-band activity tracks the content of spatial working memory”. In: Journal of Neurophysiology 115.1, pp. 168–177. DOI: 10.1152/jn.00860.2015.

Frauscher, B et al. (2018). “Atlas of the normal intracranial electroencephalogram: neurophysiological awake activity in different cortical areas”. In: Brain 141.4, pp. 1130–1144.

Freschl, J, Azizi, LA, Balboa, L, Kaldy, Z, and Blaser, E (2022). “The development of peak alpha frequency from infancy to adolescence and its role in visual temporal processing: A meta-analysis”. In: Developmental Cognitive Neuroscience 57, p. 101146. DOI: 10.1016/j.dcn.2022.101146.

Gabard-Durnam, LJ, Wilkinson, C, Kapur, K, Tager-Flusberg, H, Levin, AR, and Nelson, CA (2019). “Longitudinal EEG power in the first postnatal year differentiates autism outcomes”. In: Nature communications 10.1, p. 4188.

García-Rosales, F, Schaworonkow, N, and Hechavarria, JC (2024). “Oscillatory waveform shape and temporal spike correlations differ across bat frontal and auditory cortex”. In: Journal of Neuroscience. DOI: 10.1523/JNEUROSCI.1236-23.2023.

Gastaut, HJ and Bert, J (1954). “EEG changes during cinematographic presentation (Moving picture activation of the EEG)”. In: Electroencephalography and Clinical Neurophysiology 6, pp. 433–444. DOI: 10.1016/0013-4694(54)90058-9.

Gathercole, SE, Pickering, SJ, Ambridge, B, and Wearing, H (2004). “The structure of working memory from 4 to 15 years of age”. In: Developmental Psychology 40.2, pp. 177–190. DOI: 10.1037/0012-1649.40.2.177.

Giehl, J and Siegel, M (2024). Spectral waveform analysis dissociates human cortical alpha rhythms. DOI: 10.1101/2024.03.16.585296.

Glasser, MF et al. (2016). “A multi-modal parcellation of human cerebral cortex”. In: Nature 536.7615, pp. 171–178.

Gramfort, A et al. (2013). “MEG and EEG data analysis with MNE-Python”. In: Frontiers in Neuroscience 7. DOI: 10.3389/fnins.2013.00267.

Halgren, M et al. (2019). “The generation and propagation of the human alpha rhythm”. In: Proceedings of the National Academy of Sciences 116.47, pp. 23772–23782. DOI: 10.1073/pnas.1913092116.

Happé, F and Ronald, A (2008). “The ‘fractionable autism triad’: a review of evidence from behavioural, genetic, cognitive and neural research”. In: Neuropsychology review 18, pp. 287– 304.

Hill, AT, Clark, GM, Bigelow, FJ, Lum, JA, and Enticott, PG (2022). “Periodic and aperiodic neural activity displays age-dependent changes across early-to-middle childhood”. en. In: Developmental Cognitive Neuroscience 54. DOI: 10.1016/j.dcn.2022.101076.

Jansen, BH and Rit, VG (1995). “Electroencephalogram and visual evoked potential generation in a mathematical model of coupled cortical columns”. In: Biol. Cybern. 73, pp. 357–366.

Jarosz, AF and Wiley, J (2014). “What are the odds? A practical guide to computing and reporting Bayes factors”. In: The Journal of Problem Solving 7.1, p. 2.

Javitt, DC et al. (2020). “A roadmap for development of neuro-oscillations as translational biomarkers for treatment development in neuropsychopharmacology”. In: Neuropsychopharmacology 45.9, pp. 1411–1422. DOI: 10.1038/s41386-020-0697-9.

Jeffreys, H (1998). The theory of probability. OuP Oxford.

Jokisch, D and Jensen, O (2007). “Modulation of gamma and alpha activity during a working memory task engaging the dorsal or ventral stream”. In: The Journal of Neuroscience: The Official Journal of the Society for Neuroscience 27.12, pp. 3244–3251. DOI: 10.1523/JNEUROSCI.5399-06.2007.

Jones, SR (2016). “When brain rhythms aren’t ‘rhythmic’: implication for their mechanisms and meaning”. In: Current Opinion in Neurobiology 40, pp. 72–80. DOI: 10.1016/j.conb.2016.06.010.

Kelly, SP, Lalor, EC, Reilly, RB, and Foxe, JJ (2006). “Increases in Alpha Oscillatory Power Reflect an Active Retinotopic Mechanism for Distracter Suppression During Sustained Visuospatial Attention”. In: Journal of Neurophysiology 95.6, pp. 3844–3851. DOI: 10.1152/jn.01234.2005.

Klimesch, W (2012). “Alpha-band oscillations, attention, and controlled access to stored information”. In: Trends in Cognitive Sciences 16.12, pp. 606–617. DOI: 10.1016/j.tics.2012.10.007.

Koshino, Y and Niedermeyer, E (1975). “The clinical significance of small sharp spikes in the electroencephalogram”. In: Clinical Electroencephalography 6.3, pp. 131–140.

Kuhlman, WN (1978). “Functional topography of the human mu rhythm”. In: Electroencephalography and Clinical Neurophysiology 44.1, pp. 83–93. DOI: 10.1016/0013-4694(78)90107-4.

Kwon, H, Reiss, AL, and Menon, V (2002). “Neural basis of protracted developmental changes in visuo-spatial working memory”. In: Proceedings of the National Academy of Sciences of the United States of America 99.20, pp. 13336–13341. DOI: 10.1073/pnas.162486399.

Larson, E et al. (2022). MNE-Python. Version 1.1.0. DOI: 10.5281/zenodo.6958764.

Lefebvre, A et al. (2018). “Alpha waves as a neuromarker of autism spectrum disorder: the challenge of reproducibility and heterogeneity”. In: Frontiers in neuroscience 12, p. 662.

Lindsley, DB (1939). “A Longitudinal Study of the Occipital Alpha Rhythm in Normal Children: Frequency and Amplitude Standards”. In: The Pedagogical Seminary and Journal of Genetic Psychology 55.1, pp. 197–213. DOI: 10.1080/08856559.1939.10533190.

Lopes Da Silva, F, Van Rotterdam, A, Barts, P, Van Heusden, E, and Burr, W (1976). “Models of Neuronal Populations: The Basic Mechanisms of Rhythmicity”. In: Progress in Brain Research. Vol. 45. Elsevier, pp. 281–308. ISBN: 978-0-444-41457-1. DOI: 10.1016/S0079-6123(08)60995-4.

Lopez, K, Monachino, A, Vincent, K, Peck, F, and Gabard-Durnam, L (2023). “Stability, change, and reliable individual differences in electroencephalography measures: a lifespan perspective on progress and opportunities”. In: NeuroImage, p. 120116.

Lord, C, Elsabbagh, M, Baird, G, and Veenstra-Vanderweele, J (2018). “Autism spectrum disorder”. In: The lancet 392.10146, pp. 508–520.

Luna, B, Garver, KE, Urban, TA, Lazar, NA, and Sweeney, JA (2004). “Maturation of cognitive processes from late childhood to adulthood”. In: Child Development 75.5, pp. 1357–1372. DOI: 10.1111/j.1467-8624.2004.00745.x.

Lundqvist, M, Rose, J, Herman, P, Brincat, SL, Buschman, TJ, and Miller, EK (2016). “Gamma and Beta Bursts Underlie Working Memory”. In: Neuron 90.1, pp. 152–164. DOI: 10.1016/j.neuron.2016.02.028.

Maenner, MJ (2023). “Prevalence and characteristics of autism spectrum disorder among children aged 8 years—Autism and Developmental Disabilities Monitoring Network, 11 sites, United States, 2020”. In: MMWR. Surveillance Summaries 72.

Nigg, JT, Tannock, R, and Rohde, LA (2010). “What is to be the fate of ADHD subtypes? An introduction to the special section on research on the ADHD subtypes and implications for the DSM–V”. In: Journal of Clinical Child & Adolescent Psychology 39.6, pp. 723–725.

Nikulin, VV, Nolte, G, and Curio, G (2011). “A novel method for reliable and fast extraction of neuronal EEG/MEG oscillations on the basis of spatio-spectral decomposition”. In: NeuroImage 55.4, pp. 1528–1535. DOI: 10.1016/j.neuroimage.2011.01.057.

Oberman, LM, Hubbard, EM, McCleery, JP, Altschuler, EL, Ramachandran, VS, and Pineda, JA (2005). “EEG evidence for mirror neuron dysfunction in autism spectrum disorders”. In: Cognitive Brain Research 24.2, pp. 190–198. DOI: 10.1016/j.cogbrainres.2005.01.014.

Pfurtscheller, G and Neuper, C (1994). “Event-related synchronization of mu rhythm in the EEG over the cortical hand area in man”. In: Neuroscience Letters 174.1, pp. 93–96. DOI: 10.1016/0304-3940(94)90127-9.

Rouder, JN, Speckman, PL, Sun, D, Morey, RD, and Iverson, G (2009). “Bayesian t tests for accepting and rejecting the null hypothesis”. In: Psychonomic bulletin & review 16, pp. 225–237.

Rueda, MR, Posner, MI, and Rothbart, MK (2005). “The development of executive attention: contributions to the emergence of self-regulation”. In: Developmental Neuropsychology 28.2, pp. 573–594. DOI: 10.1207/s15326942dn2802_2.

Salmelin, R and Hari, R (1994). “Spatiotemporal characteristics of sensorimotor neuromagnetic rhythms related to thumb movement”. In: Neuroscience 60.2, pp. 537–550. DOI: 10.1016/0306-4522(94)90263-1.

Schaworonkow, N and Nikulin, VV (2019). “Spatial neuronal synchronization and the waveform of oscillations: Implications for EEG and MEG”. In: PLOS Computational Biology 15.5, e1007055. ISSN: 1553-7358. DOI: 10.1371/journal.pcbi.1007055.

Schaworonkow, N and Nikulin, VV (2022). “Is sensor space analysis good enough? Spatial patterns as a tool for assessing spatial mixing of EEG/MEG rhythms”. In: NeuroImage 253. DOI: 10.1016/j.neuroimage.2022.119093..

Schaworonkow, N and Voytek, B (2021a). “Enhancing oscillations in intracranial electrophysiological recordings with data-driven spatial filters”. In: PLOS Computational Biology 17.8. Ed. by D Marinazzo, e1009298. DOI: 10.1371/journal.pcbi.1009298.

Schaworonkow, N and Voytek, B (2021b). “Longitudinal changes in aperiodic and periodic activity in electrophysiological recordings in the first seven months of life”. In: Developmental Cognitive Neuroscience 47, p. 100895. DOI: 10.1016/j.dcn.2020.100895.

Sherman, MA et al. (2016). “Neural mechanisms of transient neocortical beta rhythms: Converging evidence from humans, computational modeling, monkeys, and mice”. In: Proceedings of the National Academy of Sciences 113.33, E4885–E4894. DOI: 10.1073/pnas.1604135113.

Snyder, AC and Foxe, JJ (2010). “Anticipatory Attentional Suppression of Visual Features Indexed by Oscillatory Alpha-Band Power Increases:A High-Density Electrical Mapping Study”. In: Journal of Neuroscience 30.11, pp. 4024–4032. DOI: 10.1523/JNEUROSCI.5684-09.2010.

Sokoliuk, R et al. (2019). “Two Spatially Distinct Posterior Alpha Sources Fulfill Different Functional Roles in Attention”. In: The Journal of Neuroscience 39.36, pp. 7183–7194. DOI: 10.1523/JNEUROSCI.1993-18.2019.

Sonuga-Barke, EJ and Halperin, JM (2010). “Developmental phenotypes and causal pathways in attention deficit/hyperactivity disorder: potential targets for early intervention?” In: Journal of Child Psychology and Psychiatry 51.4, pp. 368–389.

Stam, C, Pijn, J, Suffczynski, P, and Lopes da Silva, F (1999). “Dynamics of the human alpha rhythm: evidence for non-linearity?” In: Clinical Neurophysiology 110.10, pp. 1801–1813. DOI: 10.1016/S1388-2457(99)00099-1.

Ter Huurne, N. Lozano-Soldevilla, D, Onnink, M, Kan, C, Buitelaar, J, and Jensen, O (2017). “Diminished modulation of preparatory sensorimotor mu rhythm predicts attention-deficit/hyperactivity disorder severity”. In: Psychological Medicine 47.11, pp. 1947–1956.

Thut, G, Nietzel, A, Brandt, SA, and Pascual-Leone, A (2006). “Alpha-Band Electroencephalo-graphic Activity over Occipital Cortex Indexes Visuospatial Attention Bias and Predicts Visual Target Detection”. In: Journal of Neuroscience 26.37, pp. 9494–9502. DOI: 10.1523/JNEUROSCI.0875-06.2006.

Tröndle, M, Popov, T, Dziemian, S, and Langer, N (2022). “Decomposing the role of alpha oscillations during brain maturation”. In: eLife 11, e77571. DOI: 10.7554/eLife.77571.

Vallat, R (2018). “Pingouin: statistics in Python”. In: Journal of Open Source Software 3.31, p. 1026. DOI: 10.21105/joss.01026.

Vollebregt, MA, Zumer, JM, Ter Huurne, N. Buitelaar, JK, and Jensen, O (2016). “Posterior alpha oscillations reflect attentional problems in boys with Attention Deficit Hyperactivity Disorder”. In: Clinical Neurophysiology 127.5, pp. 2182–2191. DOI: 10.1016/j.clinph.2016.01.021.

Wetzels, R, Matzke, D, Lee, MD, Rouder, JN, Iverson, GJ, and Wagenmakers, EJ (2011). “Statistical evidence in experimental psychology: An empirical comparison using 855 t tests”. In: Perspectives on Psychological Science 6.3, pp. 291–298.

Wilkinson, CL et al. (2024). “Developmental trajectories of EEG aperiodic and periodic components in children 2–44 months of age”. en. In: Nature Communications 15.1, p. 5788. DOI: 10.1038/s41467-024-50204-4.

Worden, MS, Foxe, JJ, Wang, N, and Simpson, GV (2000). “Anticipatory Biasing of Visuospatial Attention Indexed by Retinotopically Specific α-Bank Electroencephalography Increases over Occipital Cortex”. In: The Journal of Neuroscience 20.6, RC63–RC63. DOI: 10.1523/JNEUROSCI.20-06-j0002.2000.

